# Cell-Type-Specific Surfaceome Profiling of 100-500 Isolated Cells using a Droplet-Based Magnetic Affinity Purification System

**DOI:** 10.1101/2025.03.13.643164

**Authors:** Amanda C. Lorentzian, Juliet M. Bartleson, Terence Ho, Zhichang Yang, Meena Choi, Tommy K. Cheung, Christopher M. Rose, William T. Yewdell, Ying Zhu

## Abstract

Cell surface proteins (CSPs) represent an important source of biomarkers and therapeutic targets. However, due to the inherent sensitivity limitations of existing technologies, tissue and cell-type-specific surfaceomes remain poorly characterized, especially in the context of human diseases. Herein, we develop nanoMAPS (nanoscale Magnetic Affinity Purification System), a miniaturized proteomic sample preparation method for surfaceome profiling of as few as 100-500 cells (1000× to 100,000× lower than existing technologies). We demonstrate that the miniaturization of magnetic bead-based affinity purification inside a single droplet can efficiently improve the recovery of surface proteins and reduce non-specific absorption of intracellular proteins. By applying nanoMAPS to human immune cells isolated from PBMCs, we demonstrate robust identification of both well-known cell-type-specific surface markers and candidate proteins. We establish nanoMAPS as a promising platform to expand surface proteomics from cultured cells to primary cells isolated from patients or mouse models.

## INTRODUCTION

Cell surface proteins (CSPs) are anchored or embedded in the plasma membrane and exposed to the extracellular space. Although CSPs are encoded by only ∼25% of the human genome^1^, CSPs play essential roles in many biological processes, including intercellular communications, transport regulation across the plasma membrane, antigen recognition, and the governance of cell interactions with the surrounding microenvironment^2^. Moreover, CSPs are unique to specific cell types and subtypes, which can be used to distinguish aberrant cells present in disease states^3–6^. It is increasingly evident that many diseases develop and persist through intricate interactions between highly heterogeneous cell populations and their surrounding microenvironment^7,8^. Profiling the cell surface proteome (hereafter the surfaceome) of these subpopulations of cells in disease states not only enhances our understanding of the underlying mechanisms of disease, but also holds significant promise for identifying easily accessible therapeutic targets and biomarkers^6,9^.

Tissue and cell-type-specific surfaceome profiling is essential to comprehend how specific populations of cells contribute to the underlying disease biology, and to discover therapeutic targets that are disease-specific. Compared with intracellular proteins like enzymes and transcription factors, surface proteins have distinct advantages to use as therapeutic targets. Their location on cell membrane makes them easily accessible, allowing for the use of a variety of therapeutic modalities, including small molecules, antibodies, and cell therapies. Surface proteins are often highly specific to different cell types or disease states, which can enhance the effectiveness of targeted therapies while minimizing the risk of off-target effects and toxicity. Mass spectrometry (MS)-based proteomics has been demonstrated to be an unbiased discovery technology for the cell surfaceome. However, because the abundance of surface proteins only represents < 1% of the total proteome, the technology requires a substantial amount of input material (>1 mg of protein lysate or 10 million mammalian cells), prohibiting analysis of small or rare populations of primary cells isolated from patient specimens or disease mouse models. The analysis of the surfaceome is further complicated by other technical challenges, such as lower abundance compared to their intracellular counterparts, high hydrophobicity, and few available tryptic cleavage sites^2,9^. Although recent advances in sample preparation and sensitive mass spectrometers have enabled reliable analysis with as few as 100,000 starting cells^10^, this amount is still prohibitively high for many applications using primary cells, especially given that only a limited amount of biopsy specimen can be collected from patients and multiple replicates are required to reach statistical confidence in target or biomarker discovery studies.

We aimed to develop a highly sensitive proteomics technology to enable reliable surfaceome profiling of nanogram protein quantities from <500 cells. Recent significant improvements in MS sensitivity have enabled the detection of thousands of proteins from a single cell^11,12^. Therefore, the remaining technical gap lies in the inefficient sample preparation of existing surface proteomics workflows. Compared with global proteomics workflows, surface proteomics requires the enrichment of chemically labeled proteins using magnetic beads or affinity columns^13^, which are prone to sample loss for low-input samples. Toward this end, we developed nanoMAPS (nanoscale Magnetic Affinity Purification System), a miniaturized method to perform affinity-based enrichment of nanogram quantities of biotin-labeled surface proteins using streptavidin-bound magnetic beads. NanoMAPS was developed based on the microPOTS platform^14^, a global proteomic sample preparation technology with significantly increased sample recovery for low-input samples by minimizing the assay volumes to microliter scale and performing all steps in a single droplet. Similarly, the entire nanoMAPS workflow, including cell isolation, lysis, protein binding, washing, as well as protein reduction, alkylation, and digestion, takes place in a single droplet containing low microgram beads to maximize the sample recovery and overall sensitivity.

We demonstrate that nanoMAPS not only allows for high-sensitivity surfaceome profiling of as few as 100 cells, but also facilitates the seamless integration with fluorescence-activated cell sorting (FACS) for the analysis of subpopulations from primary cells, such as human peripheral blood mononuclear cells (PBMCs). The droplet-based nanoMAPS system allows for high-efficiency CSP enrichment from intact cells, identifying 166 and 216 Surfy^15^ -annotated surface proteins from 100 and 500 HeLa cells, respectively. In applying the system to primary immune cells isolated from human PBMCs, the unbiased surface proteomics data reveals many previously validated and additional surface markers. Lastly, we demonstrate that nanoMAPS-based surface proteomics could provide more confident identification of cell type-specific CSPs compared to global proteomics.

## RESULTS

### NanoMAPS integrates surface proteomics workflow in a single droplet

**Figure 1** illustrates the overall workflow of the nanoMAPS-based surface proteomics. The sample preparation procedures were performed on a selectively patterned glass slide with an array of 2-mm-diameter microwells^16,17^. The differentially hydrophobic patterning of microwells confines microliter droplets inside the microwells during multi-step processing. The open-space microwell chip can be placed inside a FACS system for direct cell collection. Using 2-mm wells, we can collect as many as 2000 small-size immune cells (5-10 µm diameter) or 500 large-size tumor cells (15-20 µm diameter), and as few as a single cell. In a typical surface proteomics workflow, cell mixtures are initially labeled with a custom multicolor immunofluorescence panel, followed by biotin labeling (**Figure 1A**). Next, specific populations of cells are sorted via FACS directly into the 48-well nanoMAPS chip. After cell lysis and protein extraction, nanogram amounts of magnetic beads with streptavidin conjugation are added into each microwell to allow surface protein enrichment on the beads (**Figure 1B**). The magnetic beads can disperse uniformly within the microdroplet without the need for vigorous mixing, ensuring efficient protein binding (**Supplementary Figure S1)**. By applying a magnet array to the back of the microwells, the beads are quickly immobilized on the well surface, facilitating multiple-step bead washing (**Supplementary Figure S1, Supplementary Figure S2A**). Because the microwell array is spaced by 4.5 mm between two adjacent wells, the droplet array can be aligned with 384-well plates preloaded with 61 µL washing buffer to remove non-bound proteins. Because the volume ratio between the washing buffer and the droplet is ∼20:1, five wash cycles can theoretically reduce the background by >10 billion-fold. As an optional step, we transfer the protein-bound beads to a second nanoMAPS chip to further minimize potential non-specific-bound proteins on the well surface of the first chip (**Supplementary Figure S2B)**. Finally, on-bead protein reduction, alkylation, and digestion are carried out in the same droplet to generate tryptic peptides, followed by sample cleanup with Evotip, and finally analyzed with an Evosep LC and Orbitrap MS equipped with FAIMSpro interface (**Figure 1C**). To confidently identify and quantify cell surface proteins (CSPs), we only consider proteins that are significantly enriched (adjust P-value <0.05 and fold change >2) compared to negative controls (same sample without biotin labeling) and are annotated as CSPs by the Surfy database, an in silico derived surfaceome containing 2,886 proteins^15^. As described above, all the surface proteomics steps, including protein extraction, affinity purification, and tryptic digestion, are performed inside single droplets with minimal magnetic beads. Thus, compared with traditional spin-column or tube-based magnetic beads approaches, the protein recovery can be significantly improved.

**Figure 1.**
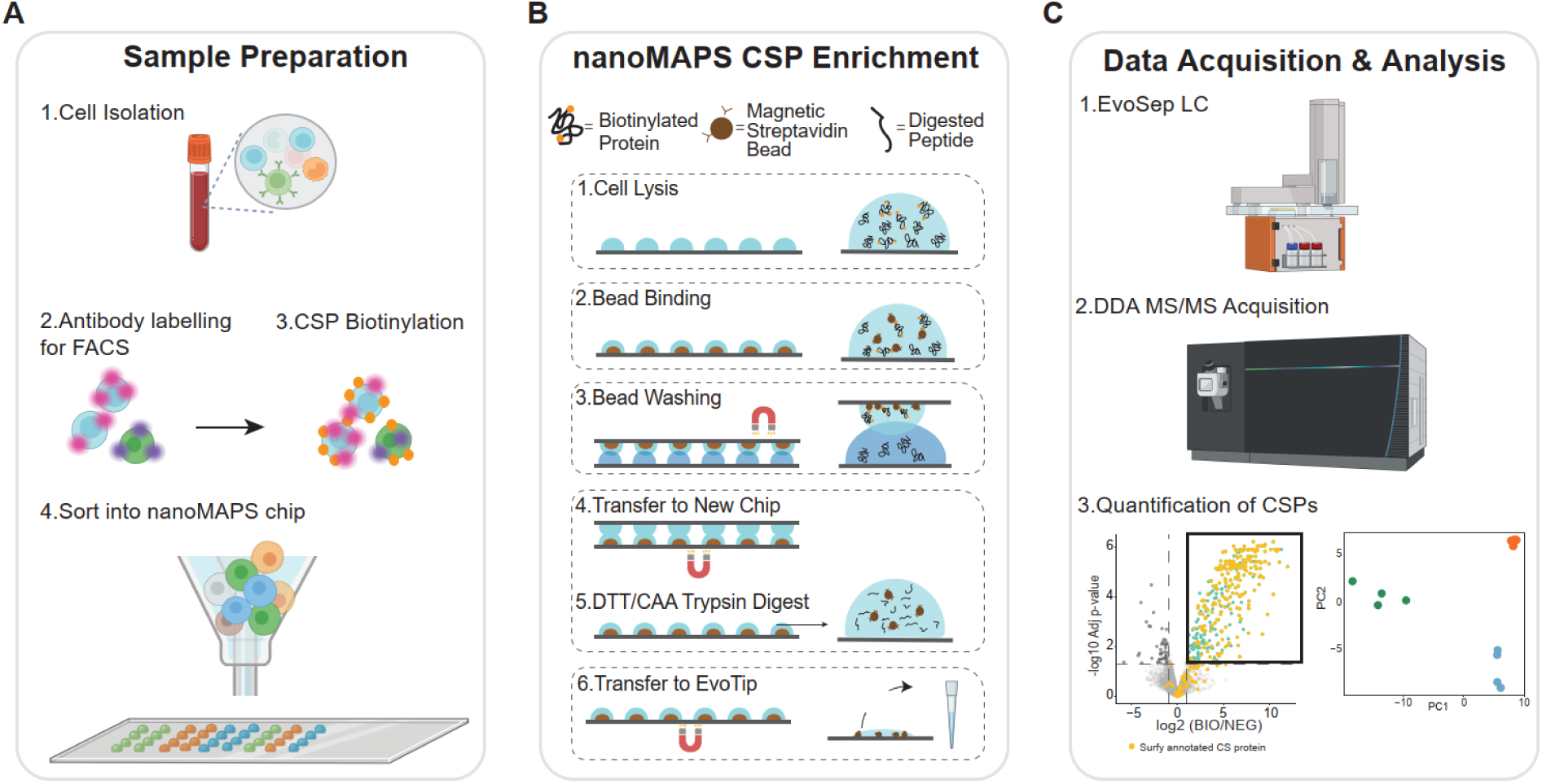
The nanoMAPS-based surface proteomics workflow for low number of FACS-isolated cells. (**A**) **Sample preparation**. Step 1. Cells are isolated from the tissue (e.g., PBMCs); Step 2. A mix of cell populations is stained with cell-type specific antibodies conjugated with fluorophores; Step 3. The cell surface proteins are biotinylated. Step 4. The cells are sorted into nanoMAPS wells by FACS. (**B**) **NanoMAPS CSP enrichment**. Step 1. The cells are lysed, and proteins are extracted inside micro droplets; Step 2. Streptavidin-bound magnetic beads are added to the droplets to isolate the biotinylated surface proteins; Step 3. The beads are rigorously washed by dipping the chip into prefilled wells of a 384-well plate to remove unbound proteins. A magnet array is attached to the back of the nanoMAPS chip to immobilize the beads during washing steps; Step 4. As an optional step, the magnetic beads are transferred to a 2nd nanoMAPS chip (chip 2) by placing a magnet below chip 2, while placing the first nanoMAPS on top of chip 2 to allow the droplet arrays to merge together; Step 5. On-bead reduction, alkylation, and trypsin digestion are performed in the same droplet; Step 6. Digested peptides are transferred and purified with EvoTips. (**C**) **Data acquisition and analysis**. Step 1. Peptides are separated with EvoSep LC; Step 2. Data is acquired on an Orbitrap MS with FAIMSpro interface; Step 3. Differential abundance analysis is used to identify the proteins that are significantly enriched in the biotinylated samples (BIO) compared to negative control samples (NEG). Cell surface protein (CSP) is confirmed if the protein is both significantly enriched and annotated as a CSP in the Surfy^15^ database. Differential abundance analysis is used to identify cell-type-specific CSPs. Figure 1 was partially created in BioRender. Lorentzian, A. (2025) https://BioRender.com/h24e838.

Since the droplet volume ranges from 1 µL to 2 µL, all these operations could be performed using a manual microliter pipette with a minimal pipetting volume of 0.2 µL. The homemade nanoMAPS chip can be replaced with a commercially available Teflon-coated microwell slide (Tekdon Inc., Myakka City, Florida) (Chip design shown in **Supplementary Figure S3**). Thus, the nanoMAPS system can readily be adopted in typical proteomics labs without significant barriers, such as the need for sophisticated devices and instrumentation. For high throughput applications, we employed a low-cost nanoliter dispenser (HP D100) for rapid reagent pipetting. We have validated that all the reagents and buffers, including the magnetic bead suspension, can be reliably dispensed into microwells with the nanoliter dispenser. It should also be noted that the droplet array format allows us to perform 48 purification assays in parallel. Thus, the throughput of nanoMAPS is on par with existing robotic magnetic purification systems, such as KingFisher from Thermo Fisher. Such high throughput is particularly useful when a large number of cell types or patient samples need to be processed for translational and biomarker discovery applications.

### NanoMAPS generates high-quality data from 1000 times less input compared to a commercial kit

We benchmarked the nanoMAPS against a commercial cell surface protein isolation kit from ThermoFisher (TF), which requires a minimum input of 1 mg of proteins. For this comparison, we first processed 1 µg of biotinylated HeLa protein lysate using both methods. NanoMAPS revealed 613 significantly enriched proteins, of which 216 (35%) are annotated as CSPs with Surfy (**Figure 2A**). Gene ontology overexpression analysis of all significantly enriched proteins confirmed their enrichment in cell surface-associated compartments, as well as in biological processes and molecular functions related to cell adhesion, receptor functions, cell communication, and integrin binding and signaling (**Figure 2B**). Although the commercial TF kit detected an average of 86 annotated CSPs from 1 µg of biotinylated HeLa lysate, the lack of reproducible data demonstrated by the low protein IDs and high coefficient of variation, prohibited us from identifying significant proteins (**Supplementary Figure S4A-C**).

**Figure 2.**
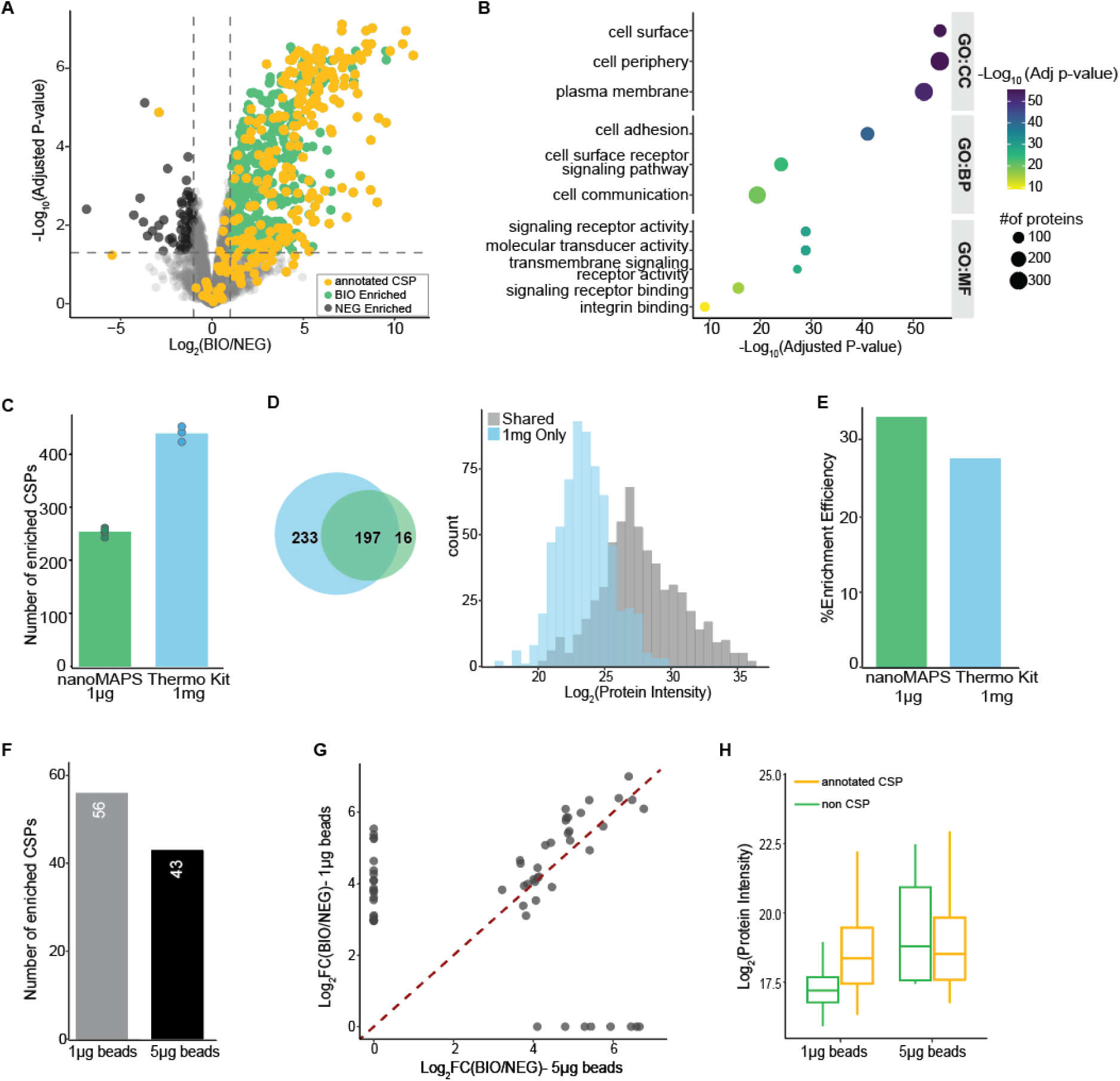
Qualitative and quantitative assessment of nanoMAPS performance in comparison with a commercial kit. **(A)** Volcano plot showing the enrichment of proteins in the biotinylated samples (BIO) compared to the negative control samples (NEG). Samples significantly enriched (Adjusted p-value <0.05 and log2FC >1, n=4) in the BIO samples are colored in green, and those annotated CSPs are in yellow. 1 µg HeLa cell lysates are used as model samples. (**B**) Gene ontology overexpression analysis using the significantly enriched proteins. Circle size represents the number of proteins matching the GO term, and color represents the –log10 (adjusted p-value). (**C**) Bar plot showing the numbers of significantly enriched CSPs from nanoMAPS-processed 1 µg HeLa cell lysates and commercial kit processed 1 mg lysate. For nanoMAPS data, four replicates (n=4) are used. For TF commercial kit, triplicates (n=3) are used. (**D**) Venn diagram showing the overlap of significant CSPs between the two workflows. The histogram indicates the distribution of the log2-transformed protein intensities for all significant CSPs from the commercial kit workflow (1-mg input). The blue bars represent proteins that are only identified in the 1 mg samples. (**E**) Bar plot showing the enrichment efficiency for the two workflows. Enrichment efficiency is calculated by dividing the number of significant CSPs by the total significantly enriched proteins. (**F**) Bar plot showing the total number of significant CSPs from nanoMAPS-processed 10-ng HeLa lysate using 1 µg and 5 µg magnetic beads (n=3). (**G**) Scatter plot of the correlation of the log2-transformed fold changes (BIO/NEG) between the nanoMAPS workflows using 1 µg and 5 µg beads. Red dashed line represents the x=y line. (**H**) Boxplot showing the distributions of the log2-transformed protein intensities of the significant proteins using 1 µg and 5 µg beads, plotted by whether a protein is annotated as a CS protein or not. Boxplot boundaries extend from the first (25%) and third (75%) quartile, representing the interquartile range (IQR). Boxplot whiskers extend to 0.75*IQR while outliers are not shown.

Next, we used the commercial TF kit to process 1 mg of HeLa lysate and compared the data quality between the two methods. Although the commercial kit used 1000 times more input, only 438 CSPs were significantly enriched (**Figure 2C**), which is ∼110% higher than the 1-µg HeLa lysate with the nanoMAPS method. Gene ontology overexpression analysis of the 438 proteins also revealed enriched categories similar to the nanoMAPS method (**Supplementary Figure S4D**). Encouragingly, 95% of significant proteins from the nanoMAPS method were included in the larger list from the commercial kit (**Figure 2D**), suggesting that both methods performed comparably in capturing CSPs. Additionally, the 233 proteins uniquely identified with the high-input TF kit protocol fall at the lower end of the intensity distribution (**Figure 2D**). This observation aligns with the expectation that these proteins are naturally low abundant proteins and would only be detected in the 1000 times higher input samples. We next investigated the proteins that were enriched in the biotinylated samples but not annotated as CSPs, and 57% of the non-CSPs enriched with the nanoMAPS workflow overlapped with those enriched in the TF workflow (**Supplementary Figure S4E**). The overlapping proteins are primarily enriched for compartments such as “extracellular vesicle”, “extracellular exosome”, and “extracellular membrane-bound organelle”, suggesting that many of these proteins that are not annotated by Surfy but enriched in our experiments are membrane associated proteins (**Supplementary Figure S4F**). However, proteins relating to the endoplasmic reticulum and cytoplasm are also enriched, demonstrating that intracellular proteins are isolated either from intracellular labelling or not thoroughly washed away. Finally, we compared the enrichment efficiency of annotated CSPs between the two workflows, which was calculated by dividing the annotated CSPs by the total significant proteins. Interestingly, 35% of all significantly enriched proteins were annotated as CSPs using the nanoMAPS method, which is slightly higher than the 27% ratio with the TF kit **(Figure 2E**). This improved ratio indicates that despite starting with 1000 times lower input, the nanoMAPS method provides comparable performance in purifying cell surface proteins while minimizing background enrichment. We hypothesize the high enrichment efficiency can be ascribed to the use of a low amount of affinity beads, as a previous study^18^ suggested an excess amount of beads can lead to the binding of non-specific background proteins, drastically reducing signal-to-noise for specific proteins.

We next investigated how bead quantity affects the enrichment efficiency of nanogram proteins. To explore this, we applied nanoMAPS to process 10-ng HeLa lysate using either 5 µg or 1 µg of magnetic beads in each droplet. Using 1-µg beads resulted in a 30% increase in identifying significant CSPs, with 56 proteins identified compared to 43 proteins using 5-µg beads (**Figure 2F**). More importantly, many of the significantly enriched CSPs using fewer beads exhibited higher log2 fold changes (log2FC) over negative control (**Figure 2G**). This can be attributed to the overall lower intensities of protein background (non-surface proteins), while comparable annotated CSP intensities between the results using 1 µg and 5 µg beads (**Figure 2H**). This result underscores the effectiveness of the nanoMAPS-based low-volume enrichment method by minimizing non-specific binding and enhancing the detection of true positives.

These data highlight the sensitivity and effectiveness of the nanoMAPS workflow in enriching cell surface proteins from low-input samples, such as 10 ng of protein lysate, which corresponds to ∼50 mammalian cells. Given that 1-µg beads exhibit improved sensitivity and specificity, we chose this amount for the following nanoMAPS experiments.

### Sensitive identification of cell surface proteins starting from 500 and 100 intact cells

Recognizing that starting from a diluted cell lysate may not accurately represent the biological conditions of intact cells, we next applied the nanoMAPS platform to an aliquot of ∼500 (average of 631 cells and 446 cells for HeLa and Jurkat respectively) or ∼100 (average of 132 cells and 97 cells for HeLa and Jurkat respectively) intact cells (**Figure 3A**). We selected HeLa cells (median diameter = 18 µm) and Jurkat (median diameter = 12 µm) cells to represent both normal-size tumor cells and small-size immune cells. We identified an average of 213 enriched CSPs from ∼500 HeLa cells (n= 4) (**Figure 3B)**. Despite only ∼100 ng protein in 500 Hela cells, the number of identified CSPs was remarkably similar to our enrichment assay starting from 1 µg of HeLa protein lysate with 81% protein overlap (**Supplementary Figure S5A**), suggesting the single-droplet nanoMAPS workflow recovers more surface proteins directly from intact cells. Encouragingly, the enrichment efficiency for the nanoMAPS assay starting from 500 intact cells was 65%, which is significantly higher compared to 33% from the previous experiment starting from cell lysate. From 100 HeLa cells (n=4), we identified an average of 166 CSPs (**Figure 3C**), and 99% of these proteins were included in the CSP list of 500 HeLa cells (**Supplementary Figure S5B**). Because Jurkat cells are only ∼30% of the volume of HeLa cells (**Figure 3D**), only 121 and 110 CSPs were identified from 500-cell and 100-cell inputs, respectively (**Figure 3E, Supplementary Figure S5C, D**). The lower CSP numbers may also be partially due to the large difference in cell functions between immune and tumor cells^5^. To further evaluate the quantitative accuracy of the CSPs in 500-cell and 100-cell inputs, we calculated the ratio of the mean protein intensities between the two inputs, expecting the ratios to be ∼5 (**Figure 3F**). Indeed, the Jurkat cells had a median ratio of 3.7, while the HeLa cells had a higher-than-expected median ratio of 7.2. The inflated ratio of HeLa cells is not well understood, but we speculate it is due to the enhanced binding of CSPs in the higher concentration of cell lysate.

**Figure 3.**
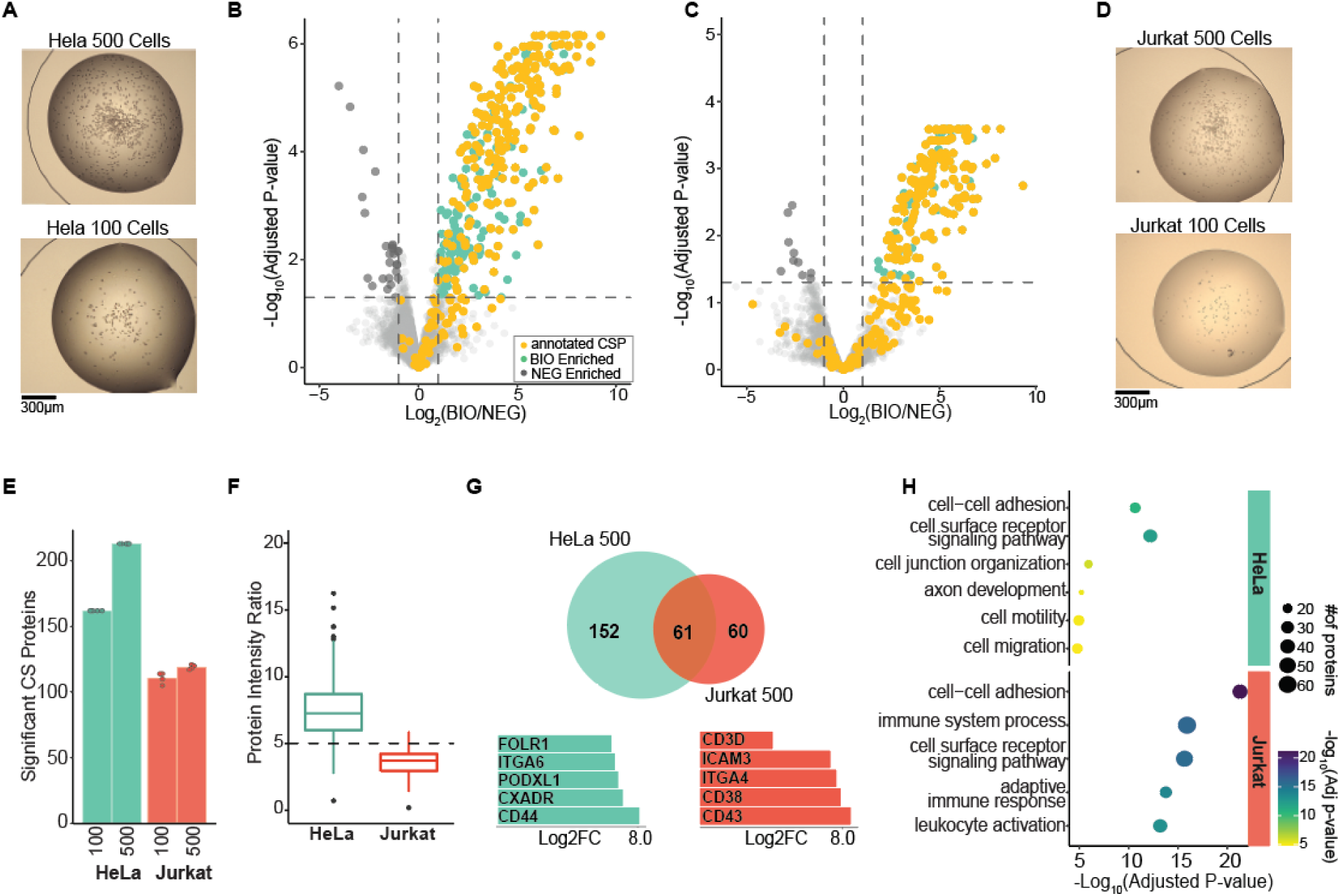
Sensitive identification of cell surface proteins starting from 500 and 100 intact cells. (**A**) Representative images of nanoMAPS wells containing ∼500 HeLa cells (Top) and ∼100 HeLa cells (bottom). (**B**) Volcano plots demonstrating the enrichment of proteins in the biotinylated samples (BIO) compared to the negative control samples (NEG) for 500 and (**C**) 100 HeLa cells. Proteins that are significantly enriched in the BIO samples (adjusted p-value <0.05 and log2FC >1, n=4) are colored in green, and those that are annotated cell surface proteins are in yellow. (**D**) Representative images of ∼500 and 100 Jurkat cells. (**E**) Barplot of significant CSPs identified in the two cell types and two cell numbers. The barplot represents the mean from four replicates. (**F**) Boxplot of the ratio of CSP intensities between the 500-cell input and the 100-cell input for HeLa and Jurkat. (**G**) (Top) Venn diagram for the overlap of significant CSPs identified from 500 HeLa cells and 500 Jurkat cells. (Bottom) Barplots showing the log2FC (BIO/NEG) of a few of the top enriched proteins in each cell type. (**H**) Gene ontology overexpression analysis of the significant CSPs in HeLa and Jurkat cells. Circle size represents the number of proteins matching the GO term, and color represents the –log10 (adjusted p-value).

Next, we evaluated whether the surfaceome profiling of the two cell populations (HeLa and Jurkat) could provide meaningful biological insights. As expected, highly distinct surface proteomes were observed between the two cell populations. Only half of the Jurkat CSPs overlapped with those identified from HeLa cells (**Figure 3G**). The top enriched CSPs from each cell type are well-defined markers and biologically relevant to their respective cell type. For HeLa cells, of notable interest was the high enrichment of cell adhesion molecules such as ITGA6, CXADR^19^, and cancer drivers and potential therapeutic targets such as FOLR1, PODXL, and CD44^20–22^. These proteins are known to be highly expressed in HeLa cells and lowly or not expressed in Jurkat cells. For Jurkat cells, important regulators of immune activity such as CD38, CD43, ITGA4, and ICAM3^23–26^, and the T-cell-specific marker CD3 were highly enriched, as expected. Finally, we performed gene ontology overexpression analysis of all the significant CSPs from both cell types. The top enriched biological process categories clearly indicated the functional differences between HeLa and Jurkat cells (**Figure 3H**). For example, the Jurkat surfaceome is enriched for processes relating to immune response and leukocyte activation, while the HeLa surfaceome is enriched for processes involving cell migration and motility, which is highly associated with tumor development and metastasis. This comparison suggests the nanoMAPS-generated cell surfaceome data is highly specific to different cell populations. It can provide unique insight into the distinct biological characteristics and functions of each cell type.

### Integration of nanoMAPS with FACS enables the identification of cell type-specific CSPs from human PBMCs

One of the main goals of mass-spectrometry-based surfaceome profiling is to identify cell-type-specific surface proteins for nominating diagnostic biomarkers and therapeutic targets. To meet this need, nanoMAPS can be directly coupled with FACS to collect different cell populations and subpopulations for high-sensitivity analysis. We evaluated the performance of nanoMAPS in the surfaceome profiling of distinct cell populations from human PBMCs, a more complex cell mixture in comparison to HeLa or Jurkat cells. Because primary cells usually contain less proteins compared with cultured cells, we collected 500 cells for each replicate and employed 4 replicates for both biotinylated cells and negative controls, requiring a total of 4000 cells for each population. This low cell number appeals to many discovery studies using primary cells from patients or disease mouse models and requires only a few hours of FACS instrument time, making the technology easily scalable for large-scale studies.

To validate the FACS-nanoMAPS workflow, we isolated three cell types from the PBMCs of a healthy donor, including monocytes (CD14+), B cells (CD19+), and T cells (CD3+) (**Figure 4A**). To minimize intracellular protein labeling, cells were biotinylated immediately after antibody labeling when the cells were most viable. 7-AAD staining is further used to gate out non-viable and permeable cells during sorting. We observed 73, 61, and 127 significant CSPs from T-Cells, B cells, and monocytes, respectively (**Figure 4B**). Because FACS enables precisely isolating the same number of cells, we evaluated the reproducibility of the nanoMAPS workflow by calculating the coefficient of variation (CV) of the enriched CSPs for each cell type (**Supplementary Figure S6)**. Although only 500 cells were processed, highly reproducible protein quantification with the median CVs of 22.9%, 26.3%, and 32.6% were obtained for monocytes, B cells, and T cells, respectively. Principal component analysis of the CSPs successfully grouped each cell type together, indicating distinct surfaceome profiles of these three cell types (**Figure 4C**). To identify cell-type-specific CSPs, we performed pair-wise differential abundance analysis and considered a protein to be cell-type specific if it was significant (adjusted p-value <0.05) and had a positive fold-change in each comparison for that cell-type. Encouragingly, many of these CSPs are uniquely enriched in each cell type with large abundance differences (**Figure 4D**). Specifically, monocytes exhibited 98 uniquely enriched CSPs, B cells had 19, and T cells had 12. The analysis identified well-established cell type-specific markers, including CD19, CD20, CD79A, and CD79B for B cells^27,28^, CD3G, CD4, CD8, and IL7R for T-cells^27^, and CD14, CD36, CD33, and CCR2 for monocytes^27,29^ **(Figure 4E**). In addition to these well-characterized surface markers, other proteins of potential interest were also detected. For instance, for monocytes, we detected markers associated with immune regulation, such as CD58 and SLAMF1, as well as key immunomodulatory proteins like CLEC12A and SIGLEC9^6^. Notably, we also identified ITGB2, a biomarker used for acute myeloid leukemia (AML) prognosis and currently being explored as a potential target for CAR-T therapy^30^. Similar activation markers were found in B cells and T cells, such as FCRL1, a potential biomarker for B cell tumors^31,32^, and BTN3A3 for T cells, which plays a crucial role in anti-tumor immune responses^33^. Though only a few were highlighted here, these data validate nanoMAPS-based surface proteomics as an unbiased approach for discovering potential biomarkers and therapeutic targets without prior knowledge or using antibodies.

**Figure 4.**
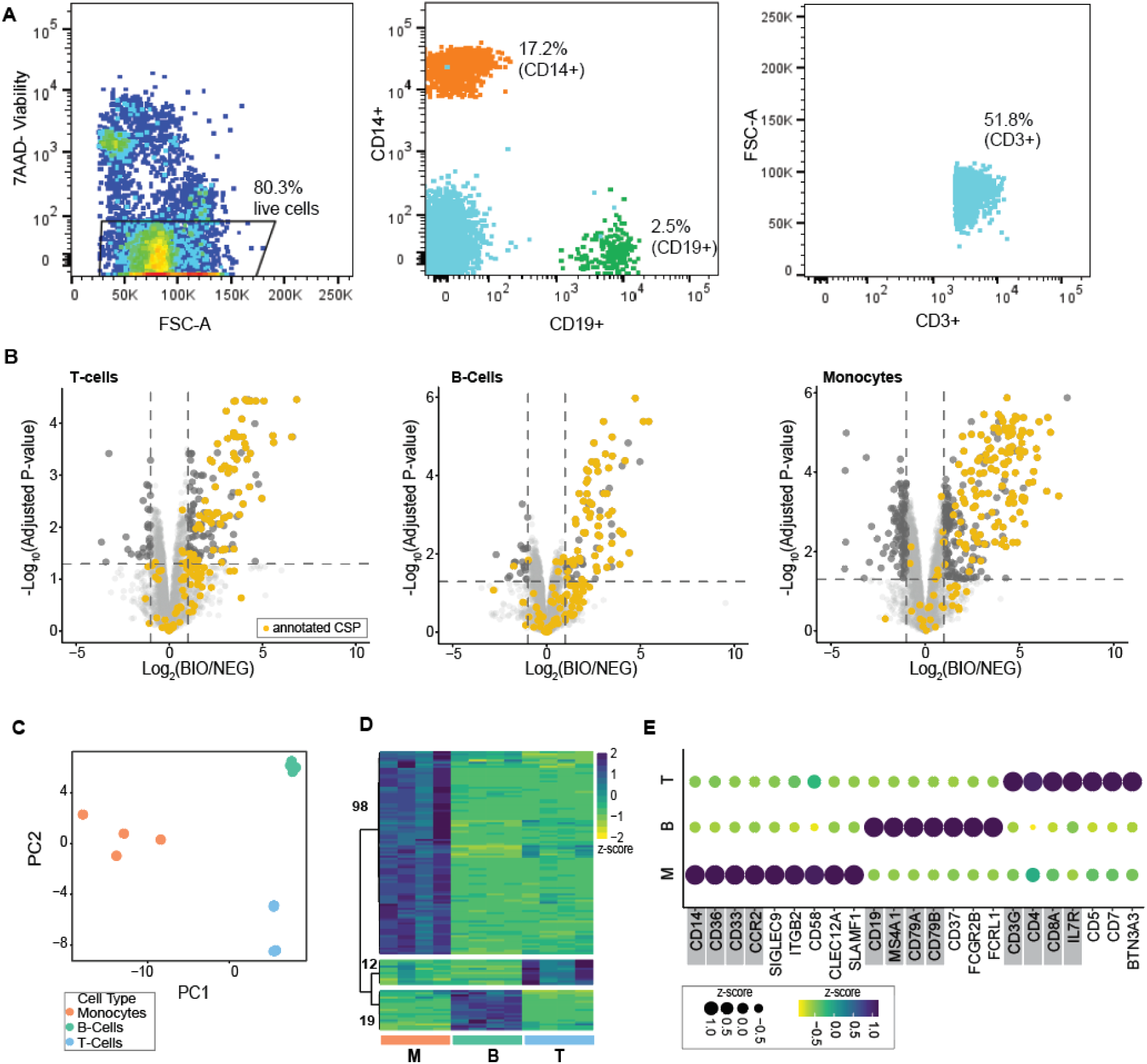
Integration of nanoMAPS with FACS enables the identification of cell-type-specific CSPs from human PBMCs. **(A**) FACS plots indicating the gating strategies to sort CD14+ monocytes, CD19+ B cells, and CD3+ T cells. Cells with low 7-Aminoactinomycin D (7-AAD) signal are gated to select only viable cells. 500 cells are collected for each replicate. (**B**) Volcano plot demonstrating the enrichment of proteins in the biotinylated samples (BIO) compared to the negative control samples (NEG) for the three cell types (adjusted p-value <0.05 and log2FC >1). Proteins that are annotated as CSPs are colored in yellow. (**C**) PCA plot showing the projection of three cell types using only significant CSPs in Figure 4B. (**D**) Pair-wise statistical analysis was used to determine cell-type-specific surface proteins. The 139 proteins with a significant enrichment (adjusted p-value <0.05) are normalized by z-score across the 12 samples and visualized in the heatmap. The numbers on the left indicate the number of proteins in each of the three main clusters. The color bars represent cell types (M for monocyte, B for B cell, and T for T cell). (**E**) A dot plot showing a selection of the most significantly distinct CSPs from the heatmap in Figure 4D. Proteins highlighted in gray are proteins that are well-characterized surface markers for the three cell types. Circle size and color represent the z-score of mean protein abundance among replicates.

Given that many studies have attempted to use transcriptomics or global proteomics to identify CSPs, we next compared the performance of nanoMAPS-based surface proteomics workflow (SP) with microPOTS-based global proteomics^16,17^ (GP) workflows in the identification and quantification of CSPs using the same number (500) of isolated cells. With the GP workflow, an average of 3,601 proteins were identified per sample (**Supplementary Figure S7A**) with CVs of annotated CSPs identified in the GP data comparable to that in the surface proteomics data (median CVs of 24.6%, 28.0%, and 25.4% for Monocytes, B Cells, and T cells, respectively) (**Supplementary Figure S7B**). The samples were also clustered by cell type, similar to the cell surface proteomics (**Supplementary Figure S7C**).

Pair-wise differential abundance analysis of all annotated CSPs revealed similar numbers of uniquely enriched CSPs (**Figure 5A**), with high overlap between SP and GP workflows for each cell type (**Figure 5B**). A notable exception was observed in T Cells, where the GP workflow identified a larger number of uniquely enriched CSPs (14 proteins) than the SP approach. Eight of these 14 proteins were not detected in the SP data. Overall, these GP-only proteins had lower abundance compared to proteins detected by both workflows (**Supplementary Figure S8A**). Another possible explanation is that these proteins are predominantly presented in intracellular locations, which is undetectable by the SP workflow. The remaining six proteins were detected in the SP data but did not pass the significance threshold for cell-type specific proteins. This suggests that these seven proteins are likely presented on the surface of other cell types as well, but the abundance ratios between intracellular and membrane locations are different (**Supplementary Figure S8B**). For example, PTPRC (CD45) and CD48, which are well-known membrane proteins for B-cells, T-cells, and monocytes^34,35^, show the highest abundance for T cells in GP data, but their abundance levels are quite similar to monocytes in SP data. Thus, the cell type differences captured by the GP data could be driven by differing levels of the intracellular counterparts of these proteins.

**Figure 5.**
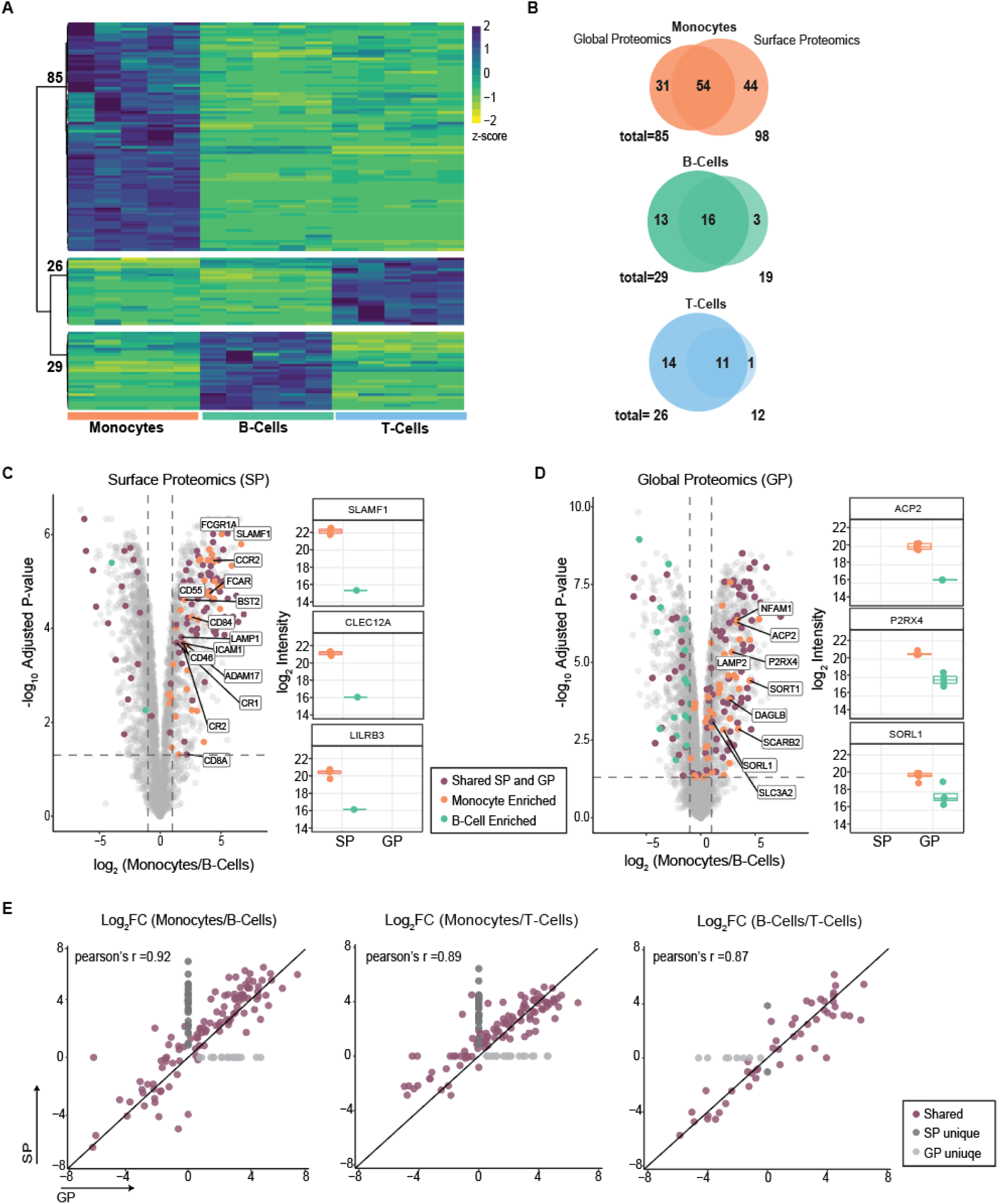
Benchmarking the performance of surface proteomics and global proteomics workflows in detecting cell-type specific CSPs. (**A**) Pair-wise differential abundance analysis was performed to identify cell-type specific cell surface proteins based on an adjusted p-value <0.05 and a positive fold change for each comparison. All significant cell-type-specific CSPs are normalized across the 15 samples by z-score and visualized in the heatmap. The numbers on the left indicate the number of proteins in each of the three main clusters. The color bars represent cell types. (**B**) Venn diagrams showing the overlap of significant CSPs from the statistical analysis for surface proteomics (SP) and global proteomics (GP). (**C**) (Left) Volcano plot demonstrating the enrichment of proteins in the monocytes compared to B cells (adjusted p-value <0.05 and log2FC >1). Proteins that were significant by both SP and GP are colored as purple. Proteins with text labels are immune response-related proteins. (Right) Boxplots of log2 protein abundance of monocyte-specific CSPs exclusively detected with SP method. (**D**) (Left) Volcano plot demonstrating the enrichment of proteins in the monocytes compared to B cells (adjusted p-value <0.05 and log2FC >1). Proteins that were significant by both SP and GP are colored as purple. Proteins with text labels are lysosome-related proteins. (Right) Boxplots to show log2 protein abundance of selected lysosome proteins from the volcano plot, which are exclusively detected in GP data. (**E**) Pair-wise correlations of log2-transformed fold-changes (log2FC) of significant CSPs measured with SP and GP. Proteins that are uniquely identified by SP are highlighted in dark grey, and proteins that are uniquely identified by GP are highlighted in light grey. R values are calculated by Pearson correlation (excluding the points that are unique to one data set).

Of particular interest, surface proteomics detected 98 uniquely enriched CSPs in monocytes, which are 13 more than the GP approach. Though the majority of proteins (54) are enriched with both workflows, 44 proteins are exclusively enriched by the SP workflow and 31 proteins are exclusively enriched by the GP workflow (**Figure 5B**). To further explore this finding, we performed statistical analysis by comparing monocytes to B cells. We chose the two cell types because they have the highest numbers of significant proteins for both surface and global proteomics based on the statistical analysis (139 proteins) (**Figure 5C, D**). Gene ontology analysis of the monocyte-enriched CSP proteins from the SP data results in enrichment terms associated with cell adhesion and immune response, which agrees well with monocyte functions **(Supplementary Figure S9**). We next focused on the proteins exclusively quantified in SP workflow and highlighted several of them in a boxplot (**Figure 5C**). Notably, these include previously mentioned monocyte CSPs, SLAMF1, a crucial immunomodulator implicated in rheumatoid arthritis^36^, and CLEC12A, a well-established prognostic marker for acute myeloid leukemia (AML)^37^, are exclusively detected in the SP workflow. Additionally, we detected LILRB3, another important immunomodulatory protein, which has also been identified as a therapeutic target for AML^38^. These clinically significant proteins were exclusively quantified by SP and were not detected by the GP approach, highlighting the unique value of the surface protein enrichment workflow to detect CSPs. Interestingly, many monocyte-enriched proteins exclusively detected in the GP workflow are associated with the lysosomal membrane and enriched to term of “protein targeting of lysosome” (**Supplementary Figure S9**), such as ACP2, SORL1, and P2RX4 (**Figure 5D**). This observation is not surprising, as monocytes have higher lysosomal activity due to their primary role as phagocytes. These data also suggest that these proteins are mainly located in lysosomes, making them more prominent in GP data and less detectable by the SP approach (**Figure 5D**). Due to their ability to traffic between the plasma membrane and lysosome during monocyte activation, these proteins are annotated as CSPs. Thus, the use of the GP approach together with Surfy annotation could potentially result in incorrectly assigned cell-type-specific CSPs, especially for proteins in multiple cellular compartments.

We further compared the quantitative performance of SP and GP workflows using significant CSPs from pair-wise statistical analysis (**Figure 5C, D**, **Supplementary Figure S10**) for all three cell types. Pair-wise correlations of log2-transformed fold-changes of the shared CSPs showed overall good agreement with Pearson correlation coefficients of > 0.85 between the two approaches (**Figure 5E**). Despite this, the fold changes were predominantly skewed towards the SP side of the x=y line, demonstrating the additional surface protein enrichment steps can better distinguish the abundance differences, due to the reduced contribution from intracellular portions of the same CSPs. Interestingly, the T cell/B cell fold changes are similar between the two methods, which can be attributed to the fact that both B and T cells are differentiated from lymphoblasts, and these significant CSPs are predominantly presented on cell surfaces. As many proteins are known to be present in both plasma membrane and intracellular compartments, surface protein enrichment allows us to differentiate the contribution of their surface portions over intracellular portions.

Together, compared with global proteomics or transcriptomics, surface proteomics can offer a more reliable and narrowed protein list for nominating potential biomarkers and targets. The ability to narrow down the protein list is highly valuable, as follow-up validation studies are time-consuming and costly.

## DISCUSSION

Surface proteins offer an invaluable resource for the discovery of diagnostic biomarkers and therapeutic targets. Indeed, ∼70% of FDA-approved drugs directly target surface proteins^39,40^. However, tissue and cell-type-specific surface proteins are still poorly characterized, especially in the context of human diseases. This is largely due to the fact that existing mass spectrometry-based proteomics technology doesn’t offer sufficient sensitivity to analyze a low number of primary cells isolated from human tissue. Herein, we validate that nanoMAPS is a highly sensitive and reliable technology for discovering cell-type-specific surface proteins using only hundreds of FACS-isolated primary cells. The technology represents a key step in expanding surface proteomics from cultured cells to primary cells with disease contexts, as hundreds of cells are readily available from patient biopsies or mouse models. Compared to transcriptomics and global proteomics, which attempt to use transcript and total protein abundance to predict surface protein abundance, the biotin-based affinity enrichment approach can minimize the contribution of intracellular proteins and offer a more confident approach to detect and quantify surface proteins. In applying nanoMAPS to cultured cells and primary PBMCs, we demonstrate hundreds of CSPs can be identified from as low as 100 cells, and highly distinct surfaceomes are presented on the surface of different cell types.

We demonstrate the miniaturization of affinity purification in a single microliter droplet with a low amount of magnetic beads greatly improves the recovery of surface proteins and minimizes the non-specific binding of background proteins. This result, together with our previous microPOTS-based global proteomics^14^ and other microfluidic affinity proteomics technologies^41,42^, highlights the critical role of reaction miniaturization for high-sensitivity proteomics. Because magnetic beads with diverse functionalizations are well-developed and validated in many bulk-scale applications, we envision nanoMAPS can become a versatile platform for low-input functional proteomics requiring affinity purification and find broad applications in studying the role of proteins in physiological and pathological contexts.

One of the key prerequisites in applying nanoMAPS for cell-type-specific surfaceome profiling is to label and sort the cell populations based on known surface markers and the corresponding antibodies. In this case, the goal of nanoMAPS assay is to use generic cell-type-specific surface markers to discover unknown ones, which are tissue or disease-specific. Such limitations can be alleviated by depleting other cell populations using an antibody cocktail and sorting the target cells using a positive/negative gating approach.

A challenging but worthy goal is the development of single-cell surface proteomics technology to obtain the surfaceome of each single cell in a complex mixture. Many technical hurdles would need to be addressed: First, the MS sensitivity needs to be significantly improved by >10 or 100 times, given the abundance of surface proteome only represents < 1% of the total proteome; Second, the sample preparation efficiency needs to be improved to enrich surface proteins from a single cell without causing significant sample losses; Third, because in many instances, the targeted cell types only account for a small percentage of total cell populations, large-scale single-cell profiling is required to cover these rare cell types. However, the low throughput of LC-MS-based proteomic technologies precludes such a large-scale study. Fortunately, recent advances^43–45^ in high-throughput proteomics have partially addressed this challenge and offered a glimpse of hope of single-cell surface proteomics. Additionally, we employed surface biotinylation coupled with negative control to minimize the impact of background proteins on CSP annotation. Such an approach doubles the number of total proteomic samples. In contrast, the cell surface capturing (CSC) technology, which involves the enrichment of cell surface N-glycoproteins and identifies CSP through tryptic peptides containing N-linked glycosylation sites, alleviates the need for negative controls^46^. Thus, the CSC method is more attractive for large-scale single-cell analysis, which we plan to evaluate in our future nanoMAPS development.

Together, we demonstrate nanoMAPS as a key step in expanding low-input global proteomics to affinity purification-based functional proteomics, where it will generate mechanistic insight into protein functions and the underlying human diseases.

## METHODS

### Cell Culture

HeLa cells were cultured at 37°C and 5% CO_2_ in Dulbecco’s Modified Eagle’s Medium supplemented with 10% fetal bovine serum and 1× penicillin-streptomycin (Sigma, St. Louis, MO, USA). Jurkat cells were cultured in RPMI 1640 supplemented with 10% fetal bovine serum and 1×penicillin-streptomycin (Sigma, St. Louis, MO, USA).

### Fabrication of nanoMAPS chip

The nanoMAPS chips are fabricated on glass microscopic slides (25.4 mm wide, 76.2 mm long) using standard photolithography and wet etching process as described previously. A total of 48 (4×12) microwells with a well diameter of 2 mm and a well-well distance of 4.5 mm were designed on a single chip. Briefly, a maskless writer (MLA 150, Heidelberg Instruments, Heidelberg, Germany) was used to transfer the design to a glass substrate with pre-coated chromium and photoresist layers (Telic Company, Valencia, CA, USA). A dose value of 90 mJ/cm^2^ and a defoc value of -2 were applied. Next, the substrate was developed with AZ® Developer 1:1 (MicroChemicals GmbH, Ulm, Germany) for 90 s, denatured at 115 ℃ for 15 min, and followed by chromium etching with Chromium Etchants 1020 (Transene, Danvers, MA, USA) for 2 min. Next, the exposed glass region was etched with buffered hydrofluoric acid (1:20) for 10 min to generate elevated pedestals (5 µm high) as microwells. The remaining photoresist was removed by soaking the substrate in acetone for 15 min with periodic shaking. The substrate was dried by placing it on a 120 °C hot plate for 1 h and then treated with oxygen plasma for 3 min (PE50XL-HF, Plasma Etch Inc., Carson City, NV, USA). The exposed glass surface was rendered to be hydrophobic by treating the substrate with 2% (v/v) heptadecafluoro-1,1,2,2-tetrahydrodecyl-dimethylchlorosilane (Thermo Scientific Chemicals, USA) in 2,2,4-trimethylpentane for 30 min. Next, the chromium layer covering the microwells was removed by chromium etchant, exposing the hydrophilic glass pedestals. To minimize the non-specific protein binding on the native glass surface, the substrate was treated with chlorotrimethylsilane vapor in a vacuum desiccator for 4 hours or overnight to passivate free silanols on the glass microwell surface.

Alternatively, the homemade nanoMAPS chips can be replaced by a commercially available Teflon-coated slide (Tekdon Inc., Myakka City, Florida, USA). We provided the design file to facilitate the adoption of the approach (**Supplementary Figure S3**). The Teflon-coated slides were gently washed with 1% (v/v) Hellmanex solution (Fisher Scientific, USA) and treated with chlorotrimethylsilane vapor to passivate glass surfaces before use.

### Biotinylation and preparation of HeLa cell lysate for comparison studies

HeLa cells adherent to the culture dish were washed twice with phosphate-buffered saline (PBS) (Sigma, St. Louis, MO, USA). Biotinylation reagent was prepared and added into the dish following the manufacturer’s protocol for Pierce Cell Surface Biotinylation and Isolation kit (ThermoFisher Scientific, Waltham, MA, USA). Cells were incubated with biotinylation reagent at room temperature (RT) for 10 minutes. Next, the biotinylation reagent was removed, and the cells were washed twice with ice-cold Tris Buffered Saline (TBS) to quench the remaining biotinylation reagent. Cells were scraped from the dish, transferred to a 15mL conical tube, and centrifuged at 300xg for 5 minutes. Cells were resuspended in either 1 mL of kit-provided lysis buffer or 1 mL of nanoMAPS lysis buffer containing 1x RIPA (ThermoFisher Scientific) with 0.1% (m/v) n-dodecyl-β-D-maltoside (DDM) (Anatrace, Maumee, OH, USA), and transferred to a 1.5mL Eppendorf tube. Additional cells for negative control were prepared in the same manner. However, the biotinylation reagent was replaced with PBS.

The lysates were sonicated using a cuphorn sonicator, alternating between 1 minute of sonication (pulsing every other second) and 1 minute on ice for 3 rounds. The lysates were then incubated on ice for 30 minutes following manufacturer’s protocol. The lysates were centrifuged at 15,000xg for 5 min at 4°C and the supernatant was transferred to a new tube. Protein concentrations were measured using Pierce Rapid Gold BCA Protein Assay Kit (ThermoFisher Scientific).

### Cell surface protein enrichment with a commercial kit

Lysate was diluted to either 1 µg or 1 mg in 500 µl of lysis buffer. Samples were prepared following the manufacturer’s protocol. Following the suggestion in the protocol, samples were de-salted using the Pierce EasyPep MS Sample Prep kit (ThermoFisher Scientific). Peptide concentrations for the samples that started with 1 mg input were measured with Pierce Quantitative Colorimetric Peptide Assay (ThermoFisher Scientific).

### Cell surface protein enrichment with nanoMAPS chips starting from HeLa lysate

Previously prepared HeLa lysate was diluted to 1 µg/µl and aliquoted directly into the microPOTS chip. Dynabeads MyOne C1 streptavidin magnetic beads (ThermoFisher Scientific) were added to each well at 10 µg/µL. The chip was covered with a plastic cover and wrapped in aluminum foil, then incubated for 2 hours at 4°C with end-over-end mixing.

48 wells of 384 well plates were filled with 61 µl per well of RIPA lysis buffer. The first nanoMAPS chip (Chip 1) was placed upside down in the magnetic holder, with the magnet on top of the chip to keep the beads held to the chip (see **Supplementary Figure S2 and Figure 1**). The holder was placed into the 384 well plate such that the micro droplets from the chip and the washing buffer in the wells were merged together. This process was repeated twice with RIPA for a total of three RIPA washes, three times with 0.5 M NaCl, two times with water to remove the salt, and one last time with 50 mM ammonium bicarbonate buffer (ABC) as a buffer exchange. For each wash, the holder is lifted up and down a few times to facilitate the mixing, and incubated for 1 minute. For the second RIPA wash and the second 0.5 M NaCl wash, Chip 1 is removed from the holder and incubated for 5 minutes with end-over-end mixing at 4°C as previously described.

As an optional step, a second nanoMAPS chip (Chip 2) was prepared with 1 µl of 5 mM dithiothreitol (DTT, No-Weigh Format, ThermoFisher Scientific) in ABC in each well and placed on top of a second magnetic holder. Next, Chip 1 was placed upside down on the second holder such that the drops from Chip 1 and Chip 2 are slightly touching in a pair-wise manner. The magnetic beads containing the bound biotinylated proteins are then pulled down from Chip 1 into Chip 2.

The chip (Chip 1 or Chip 2) containing magnetic beads is covered with a plastic cover, wrapped in foil, placed in a humidity chamber, and incubated 60°C for 15 minutes to denature and reduce the proteins. If the above optional bead transfer step is not performed, we replace the buffer in Chip 1 with 1 µl of 5 mM DTT in ABC buffer before the incubation. The humidity chamber is an empty pipette tip box with a wet paper towel in the bottom compartment, with the chip placed on top of the empty tip rack. It is then closed and placed into a Ziploc bag with a few droplets of water. The entire chamber is pre-warmed at the required temperature prior to each incubation step. Next, 0.5 µl of 30 mM chloroacetamide (CAA) (ThermoFisher Scientific) is added for a final concentration of 10 mM, and incubated at RT in the dark for 40 min. Finally, 10 ng of trypsin (Mass Spectrometry Grade, Promega, Madison, WI) in 0.2 µl ABC is added to each well. The chip is incubated in the humidity chamber for 10 hours at 37°C. Cells were acidified with 0.1% formic acid (FA) (ThermoFisher Scientific) in 0.02% DDM and transferred to a low-binding autosampler vial (PSVial100, ProteoSave, 0.3 mL, AMR Inc., Meguro-ku, Tokyo, Japan) for mass spectrometric analysis.

### Cell surface protein enrichment with nanoMAPS workflow starting from intact cells

HeLa cells were biotinylated or treated with PBS (negative control) as described above. To biotinylate Jurkat cells, they were transferred to a 15mL conical tube and centrifuged at 300xg for 5 minutes to pellet. Then the cells were washed twice with PBS. After the removal of PBS, the pellet was resuspended in 10 mL of biotinylation reagent and incubated at RT for 10 minutes. After washing twice with ice-cold TBS, cells were resuspended in PBS for cell count and viability check. Cells were diluted to either 1000 cells/µl for the 500-cell experiment or 200 cells/µl for the 100-cell experiment. Then, 0.5 µl of the cell mixture was pipetted into each well of a nanoMAPS chip. To confirm the number of cells per well, we inspected the nanoMAPS chip on a Zeiss microscope (PALM microbeam, Carl Zeiss, Jena, Germany) and took two representative images to count cells. After the evaporation of PBS, 1 µl of lysis buffer (RIPA, 0.1% DDM, and 0.5u/µl of benzonase (Sigma)) was added to each well containing the cells, then the chip was incubated in the humidity chamber at 37°C for 25 minutes. The remainder of the protocol was performed as previously described, except that 1 µg of magnetic beads instead of 5 µg were added to the wells.

### FACS isolation of peripheral blood mononuclear cells (PBMCs) into nanoMAPS and microPOTS chips

PBMCs were from a 60-year-old healthy male donor, commercially sourced from CGT Global (Folsom, CA, USA). The donor information is de-identified. The cells were thawed and washed according to the provider’s instructions. Briefly, 1 tube (∼ 15 million cells) was thawed in a 37°C water bath for 3-5 minutes, until the sample was 75% thawed. The sample was then transferred to 10mL of pre-warmed RPMI media supplemented with 10% FBS in a 15mL conical tube. The cells were centrifuged at 300xg for 5 minutes, and then washed twice with PBS. After the second wash, cell count and viability test were performed. A total of 12×10^6 cells in 1.2 mL of FACS buffer (1x PBS, 0.5% BSA, 0.05% Na Azide) was taken for staining. Cells were first incubated with Human TruStain FC-block (Biolegend) for 10 minutes at RT. Next, a master mix of the antibodies was prepared in the following manner: CD3-FITC (clone UCHT1) (Biolegend, San Diego, CA, USA) at 5µl/1×10^6 cells, CD19-PE (clone H1IB19) (Biolegend) at 5 µl/1×10^6 cells, and CD14-APC (clone M5E2) (Biolegend) at 5 µl/1×10^6 cells. To compensate for fluorescence spillover, compensation beads (1µL/1 drop of beads) were used for each fluorophore. The cells were incubated with antibodies at 4°C for 25 minutes, then washed twice with PBS. The sample was split in half to proceed with either surface biotinylation or treatment with PBS (negative control). Biotinylation was performed as previously described. After biotinylation and washing, the cells were passed through a 100 µm strainer to remove clumped cells prior to FACS. The 7AAD viability dye (Biolegend), is added just prior to sorting at a concentration of 5 µl/1×10^6 cells in 500 µl of PBS.

Sorting was performed on a BD FACSymphony S6 Cell Sorter. Sorting was conducted using 70-micron nozzles with a 70 PSI setting. These settings enable sorting of up to 2000 cells in 2 µL, the max volume of the nanoMAPS chip. BD FACS Accudrop Beads were used to calibrate sort timing, following the BD method outlined in the user manual. A custom plate setting, along with adjustments to the plate loader position and stream voltage slider, ensured the droplet deposits were aligned to the center of the wells. Single-cell sorting mode was employed to deposit cells accurately during sorting. As part of the sort-gating strategy, two singlet gates (SSC-W vs. SSC-H and FSC-W vs. FSC-H) were used to help remove doublets. Five hundred cells of each cell type were sorted directly into a nanoMAPS chip for surface proteomics, and a microPOTS chip for global proteomics. A 3D-printed holder replicating a 384 well-plate was used to hold the chips in place and enable seamless alignment with the sorter. However, the chip may also be taped to the top of the 384 well-plate and will be compatible with any instrument that can sort into a 384 well format.

### Global proteomic sample preparation from FACS-sorted Peripheral blood mononuclear cells (PBMCs) with microPOTS

Cells are lysed in 1 µl of lysis buffer (50mM ABC, 100mM NaCl, 5mM TCEP, 0.1% DDM) and incubated in the humidity chamber at 70°C for 30 minutes. Next, 0.5 µl of 30mM chloroacetamide (CAA) is added for a final concentration of 10mM, and incubated at RT in the dark for 40 min.10 ng of trypsin in 0.2 µl ABC is added to each well. The chip is incubated in the humidity chamber for 10 hours at 37°C. Cells were acidified with 0.1% FA in 0.02% DDM and transferred to EvoSep Pure Tips.

### Automated cell surface enrichment with the HP D100

A 3D-printed adapter was used to fit the nanoMAPS chip with the HP D100 (HP Inc., Palo Alto, California, USA) to make it compatible with the 384 well plate template for dispensing. The chip is loaded onto the adapter and set into place on the HP system. The 384 well plate layout is selected. The first step is to dispense 1 µl of lysis buffer (RIPA, 0.1% DDM and 0.5u/µl of benzonase) in the nanoMAPS chip containing cells. Select the “Aqueous” liquid class, and “Aqueous + Brij 35” as the buffer. Two T1 cartridges were used to dispense the volume into the chip. After cell lysis, using the same parameters, 0.5 µl of the bead mixture (1:5 dilution of beads in lysis buffer) was dispensed. Bead binding and washing were performed as described in previous steps. The following steps were performed using T1-SF cartridges with “Aqueous” selected for liquid class and “Aqueous (surfactant free)” selected for the buffer. For bead transfer to a new chip, the second nanoMAPS chip was prefilled (Chip 2) with 1 µl of 5 mM DTT in 50 mM ABC using T1-SF cartridges. The chip was incubated in the humidity chamber for 15 minutes at 60°C for protein reduction. For alkylation, 0.5 µl of 30 µM CAA in 50 mM ABC is dispensed using the same parameters as DTT. Finally, 0.2 µl of 50 ng/µl Trypsin is dispensed in the chip, then incubated in a humidity chamber for 10 hours at 37°C.

### LC-MS/MS Analysis

A Thermo Fisher UltiMate 3000 RSLC nano System was used for HeLa cell lysate and aliquoted intact cell (HeLa and Jurkat) experiments. The sample preloaded in low-binding autosampler vials (PSVial100, 0.3 mL, AMR Inc. Meguro-ku, Tokyo, Japan) was fully loaded into a 10-µL sample loop and then trapped on a capillary solid phase extraction (SPE) trap column (Acclaim™ PepMap™ 100 trap column, Thermo Fisher Scientific). Next, the peptides were separated on a 25-cm IonOpticks Aurora Series column (75 µm ID, 1.7 µm C18, IonOpticks, Melbourne, Australia) operated at 200 nL/min and a 60 min gradient from 5% to 45% Buffer B (0.1% formic acid in 98% acetonitrile). The analytical column was heated to 50 °C using Sonation Nanospray Flex column oven (Sonation GmbH, Biberach, Germany).

EvoSep LC (Evosep Biosystems, Odense, Denmark) was used for the PBMC experiment. EvoSep Pure Tips were conditioned following manufacturer’s protocol. Briefly, 20 µl of Buffer B (0.1% FA in 100 % Acetonitrile) was added to each tip and then centrifuged for 1 min at 400 x g. Next, the tips were soaked in isopropyl alcohol (IPA) enough that the disc inside the tips were fully submerged and saturated (1-2 min). Then 20 µl of Buffer A (0.1% formic acid in water) was added and centrifuged for 1 min at 400 x g. For sample loading, the tips were prefilled with 20 µl of Buffer A. Samples were transferred directly from the nanoMAPS chip to EvoSep tips, and the loaded peptides were washed once with 20 µl of Buffer A. Samples were eluted and separated using the 20SPD whisper zoom method and a 15-cm IonOpticks Aurora Elite XT column (75 µm ID, 1.7 µm C18) heated with an IonOpticks column heater (Melbourne, Australia).

An Orbitrap Eclipse Tribrid MS (ThermoFisher Scientific) with an FAIMS (high-field asymmetric-waveform ion-mobility spectrometry) interface, operated in data-dependent acquisition mode, was used for all analyses. Source settings included a spray voltage of 1,800 V, an ion transfer tube temperature of 250°C, and a FAIMS carrier gas flow of 3.5 L/min. Ionized peptides were fractionated by the FAIMS interface using two compensation voltages (-45V and -65V) with a cycle time of 1.2 s per compensation voltage. Fractionated ions within a mass range of 375-1600 m/z were acquired in Orbitrap at 120,000 resolution with a max injection time of 246 ms, an AGC target of 1E6, RF lens of 30%. Precursor ions with charge states from +2 to +6 and intensities >1E4 were collected with an isolation window of 1.4 m/z, an AGC of 2E4, and a maximum injection time of 86 ms. The isolated ions were fragmented at a high energy dissociation (HCD) of 30% and then detected in an ion trap using a “rapid” scan rate.

### Database searching and data analysis

All proteomic data raw files were processed by FragPipe (version 20.0) and searched against the *Homo sapiens* UniProt protein sequence database with decoy sequences (UP000005640 containing 20,457 forward entries, accessed 08/2023). Search settings included a precursor mass tolerance of +/-20 ppm, fragment mass tolerance of +/-0.5 Da, deisotoping, strict trypsin as the enzyme (with allowance for N-terminal semi-tryptic peptides), cysteine carbamidomethylation as a fixed modification, and several variable modifications, including oxidation of methionine, and N-terminal acetylation. Protein and peptide identifications were filtered with a false discovery rate (FDR) of less than 1% within FragPipe. IonQuant MBR (MBR) and MaxLFQ were enabled for label free quantification. An MBR FDR of 5% at ion level, a retention time tolerance of 0.4 min, a minimum of 2 isotopes, an m/z tolerance of 10 ppm, and a minimum of 3 scans were used to reduce false matching. FragPipe result files (“combined_protein.tsv”) were then imported into RStudio (Build 461) for downstream analysis in the R environment (version 4.3.1).

For surface proteomics experiments, median-centering normalization was performed in R such that each protein within a sample is divided by the median of all proteins for that sample. Then all values in the dataset are multiplied by the median of the whole dataset to correct for any scale discrepancies and bring them back to a typical magnitude range. For each experiment, all biotinylated samples were normalized as one group, and all negative control samples were normalized as one group, with the assumption that the biotinylated sample groups would have higher median intensities than negative control groups. For global proteomics, all the groups were normalized together. Prior to differential abundance analysis, the protein list was filtered such that a Protein ID remained only if it was identified in at least 50% of the replica in at least one group. Then, the missing data were imputed using Perseus-Type imputation^47^. Briefly, missing values are replaced by random numbers following a normal distribution with a standard deviation of 0.2 and downshifted by a magnitude of 2. LIMMA (package limma v 3.56.2)^48^ was used for all differential abundance analyses. For the identification of CSPs from human samples, pair-wise differential abundance analysis for each cell-type comparison was performed (monocytes vs B cells, monocytes vs T cells, and B cells vs T cells) using LIMMA. A protein was determined as cell-type specific if it had a positive fold change over all other cell types and an adjusted p-value <0.05 in each of its comparisons. For SP, that list was then filtered for validated CSPs (proteins that were enriched in the BIO/NEG comparison and annotated as CSP by Surfy). For GP, the list was only filtered for proteins annotated as CSP by Surfy. All heatmaps were created with the R package pheatmap (v1.0.12) and clustering was performed using the ward.D clustering method. Gene ontology analysis was conducted with gProfiler (version e111_eg58_p18_f463989d)^49^, using all proteins identified in the data set as the custom background. The default “user threshold” (adjusted p-value) of 0.05 was used for a significance filter.

## DATA AVAILABILITY

The mass spectrometry raw data have been deposited to the ProteomeXchange Consortium via the MassIVE partner repository with dataset identifier MSV000097303.

## ACKNOWLEDGMENTS

We thank the Genentech internal anonymous reviewers for their critical reading and suggestions of the manuscript. A.C.L. acknowledges the financial support of Genentech postdoctoral program. The work was supported by Genentech Inc.

## AUTHOR CONTRIBUTIONS

Y.Z. and A.C.L. conceptualized and designed the research. A.C.L. performed cell culture and T.H. performed the FACS sorting. A.C.L. performed microchip fabrication and sample preparation for surface proteomics and global proteomics. LC-MS analysis performed by A.C.L., Z.Y., and T.K.C. Data analysis and visualization were performed by A.C.L with insight from M.C. and Y.Z. A.C.L and Y.Z. drafted the manuscript. C.M.R. and W.T.Y. edited the manuscript. All authors have read and approved the final manuscript.

