## Supplementary information for "Cell-Type-Specific Surfaceome Profiling of 100-500 Isolated Cells using a Droplet-Based Magnetic Affinity Purification System"

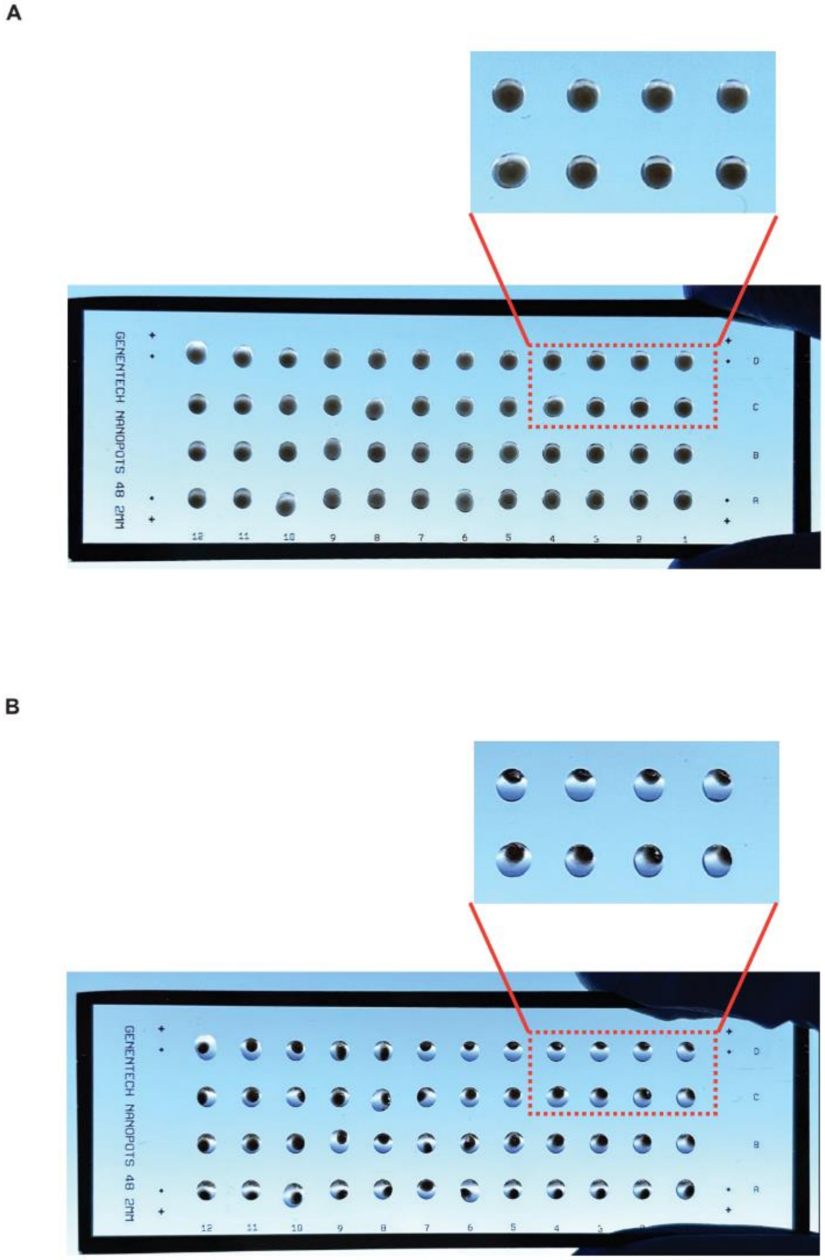

**Supplementary Figure S1**. Droplet array with magnetic beads without and with magnets. (**A**) An array of droplets containing 5 µg streptavidin magnetic beads to show the uniform bead distribution. (**B**) The magnetic beads are aggregated together when an array of magnets are applied.

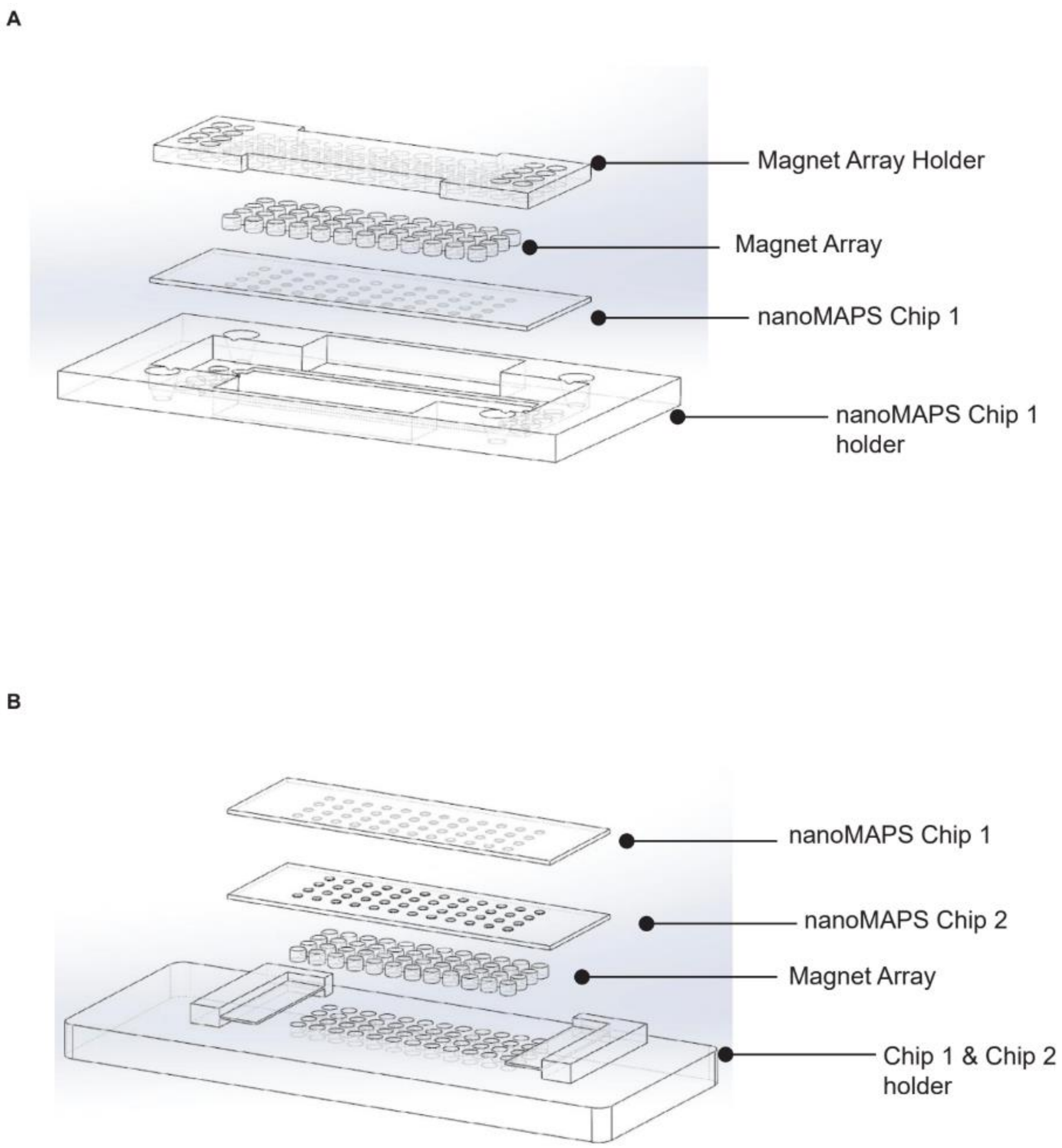

**Supplementary Figure S2**. Schematic design of 3D-printed holders for nanoMAPS chips. (**A**) Diagram of nanoMAPS chip holder 1 for bead washing steps. The device includes a holder for the chip to be placed upside down so that a magnet array can be placed on the back of nanoMAPS chip to keep the beads held to the surface of the chip while the washes occur. (**B**) Diagram of nanoMAPS child holder 2 for bead transfer from Chip 1 to Chip 2. The device has a magnet array for Chip 2 (containing buffer). There is a separation ledge for Chip 1 (containing beads) to be placed on top of, upside down, such that the droplets from each chip are slightly touching. This enables the magnetic beads to be pulled down from Chip 1, into the wells of Chip 2.

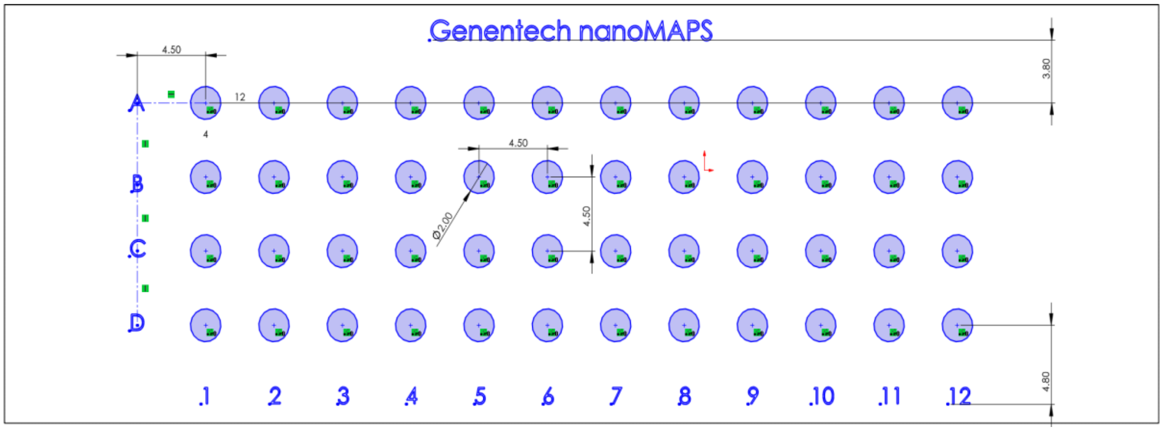

**Supplementary Figure S3**. Diagram of the 48-well nanoMAPS chip for making Teflon coated slide from Tekdon, Inc.

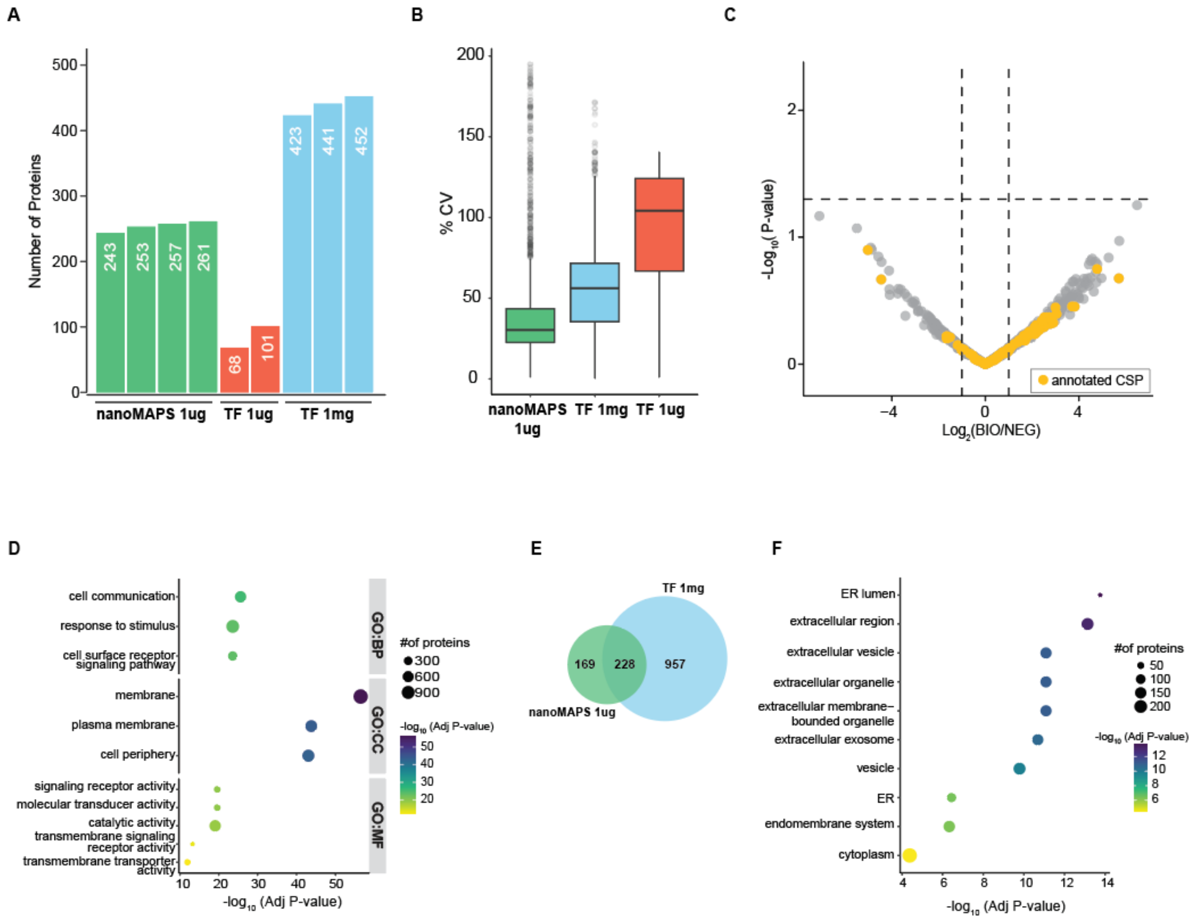

**Supplementary Figure S4**. Comparison of cell surface protein enrichment workflows between nanoMAPS and ThermoFisher kit.

(**A**)The numbers of annotated CSPs (not tested for significance between BIO and NEG) detected from nanoMAPS (1 µg input) and Thermo Fisher (TF) kit (1 µg and 1 mg inputs). (**B**) Volcano plot showing the enrichment of proteins in the biotinylated samples (BIO) compared to the negative control samples (NEG). (**C**) Coefficient of variation (cv) of protein intensity all proteins detected across the replica in each sample group. (**D**) Gene Ontology analysis for the significant CSPs from 1 mg HeLa protein lysate based on the Thermo Fisher kit. Size of the circles represent the number of proteins and the color of the circles represents the -log10 adjusted p-value of the enrichment. (**E**) Overlap of non-CSPs enriched in the biotinylated samples detected from nanoMAPS (1µg input) and TF kit (1 mg input). (**F**) Gene Ontology (cellular compartment) analysis for the 228 enriched non-CSPs that were detected by both methods. Size of the circles represent the number of proteins and the color of the circles represents the -log10 adjusted p-value of the enrichment.

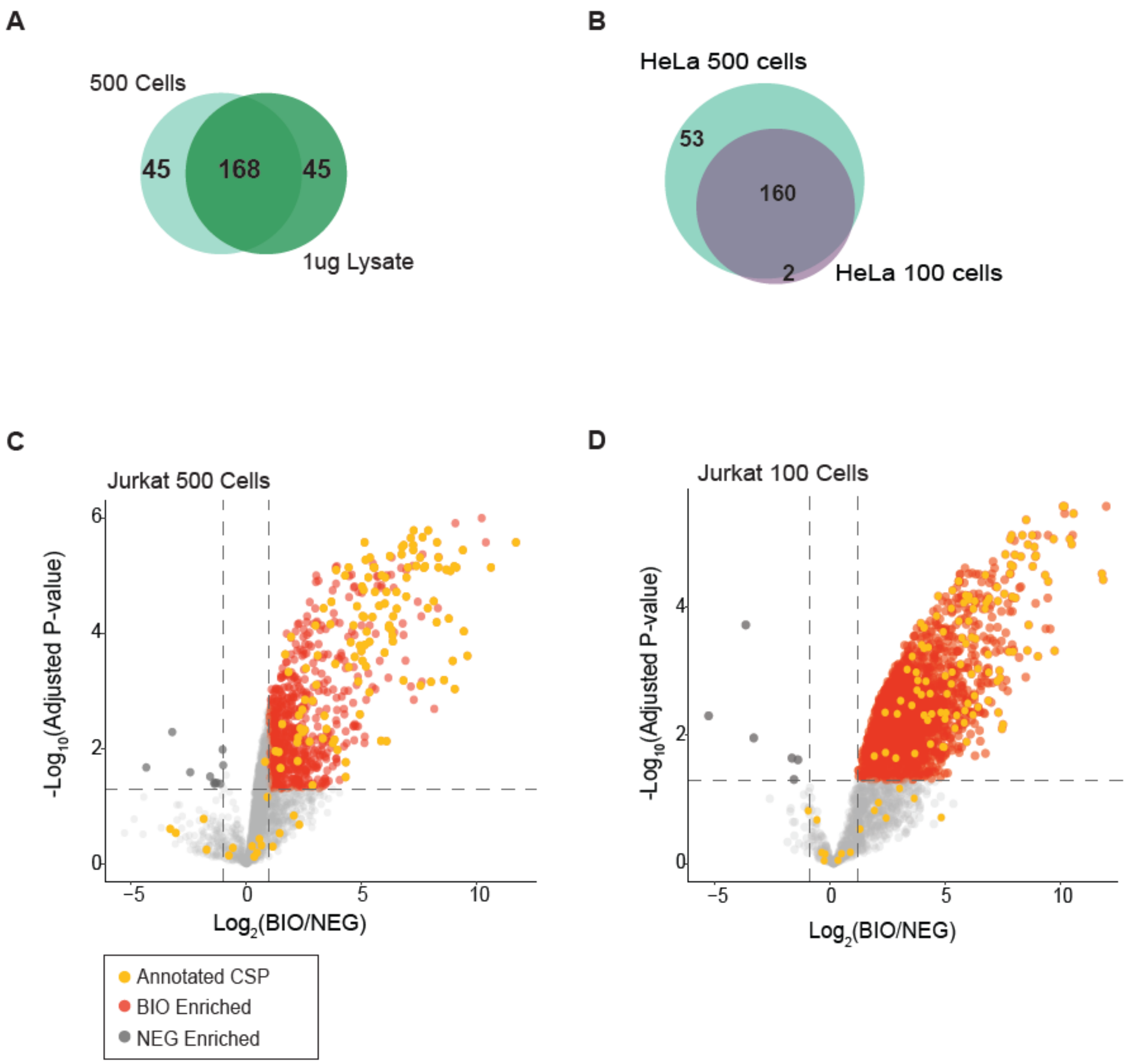

**Supplementary Figure S5**. Evaluation of nanoMAPS workflow starting from 100 and 500 intact cells.

(**A**)Venn diagram of significant CSPs identified with the nanoMAPS workflow starting from 500 HeLa cells and 1 µg HeLa lysate. (**B**) Venn diagram of significant CSPs identified with the nanoMAPS workflow starting from 500 HeLa cells and 100 HeLa cells. (**C**)Volcano plot showing the enrichment of proteins in the biotinylated samples (BIO) com- pared to the negative control samples (NEG) for starting from 500 Jurkat cells. Proteins that are annotated as CSPs are colored in yellow. (**D**) Volcano plot showing the enrichment of proteins in the biotinylated samples (BIO) compared to the negative control samples (NEG) for starting from 100 Jurkat cells. Proteins that are annotated as CSPs are colored in yellow.

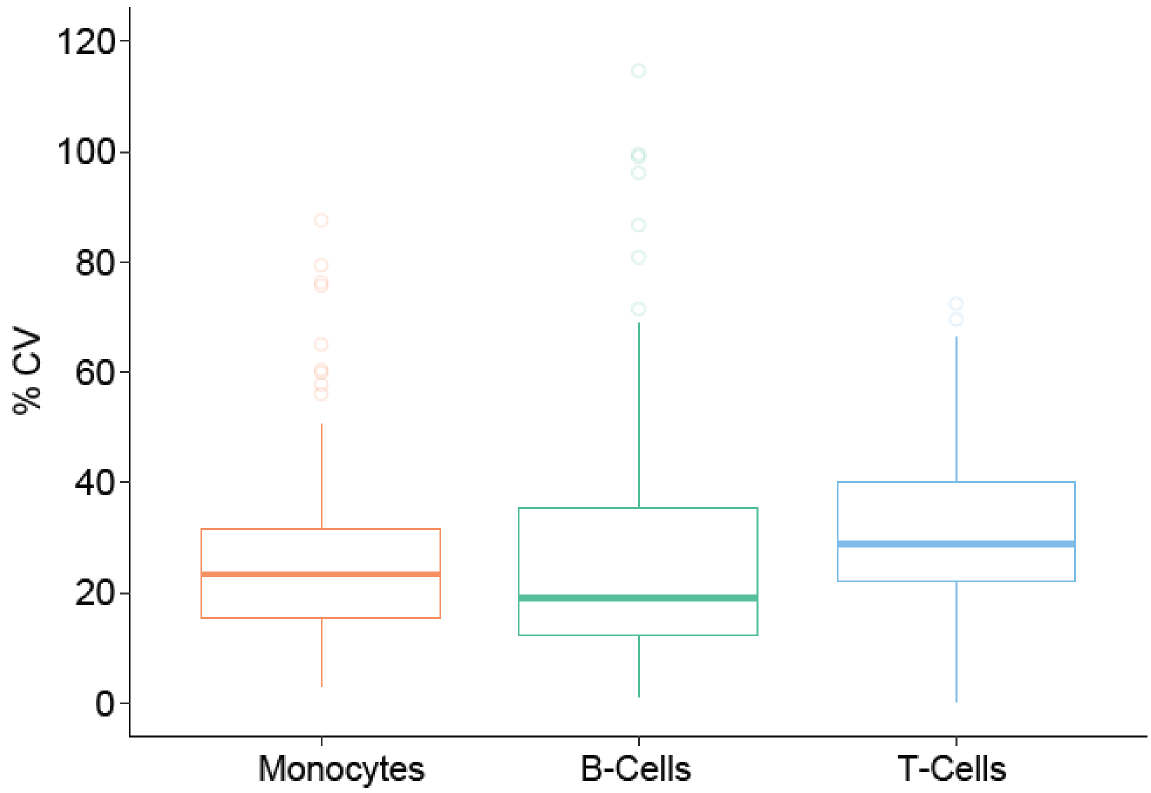

**Supplementary Figure S6**. Boxplot for CVs of significant CSPs quantified across replica (n=4) for the nanoMAPS workflow starting from 500 cells for each cell type.

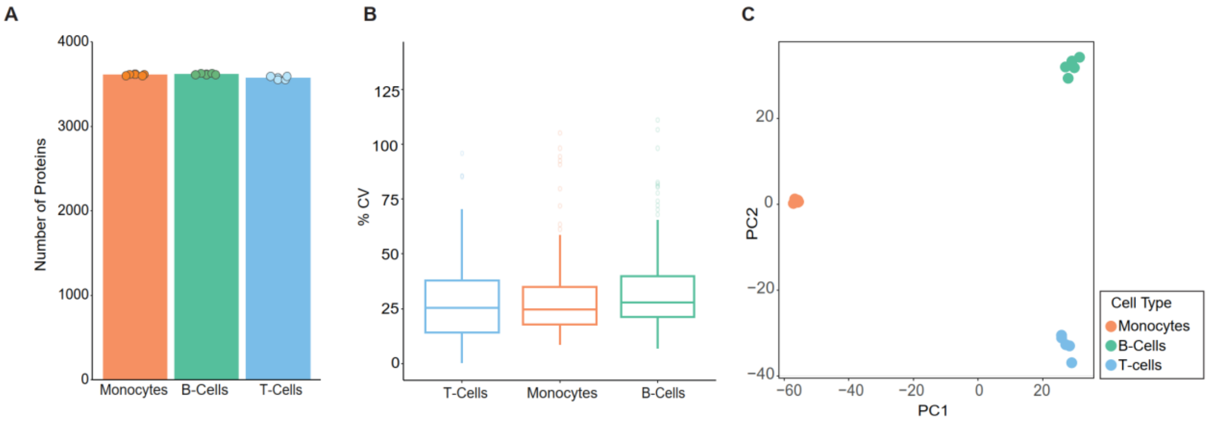

**Supplementary Figure S7**. Global proteomics of the three type of cells isolated from PBMCs. (**A**)Total number of proteins identified in each cell type with microPOTS-based global proteomics (GP). (**B**) Boxplot for CVs of annotated CSPs quantified across replica (n=5) for the GP workflow starting from 500 cells for each cell type. (**C**) PCA plot showing the grouping of three cell types using total identified proteins.

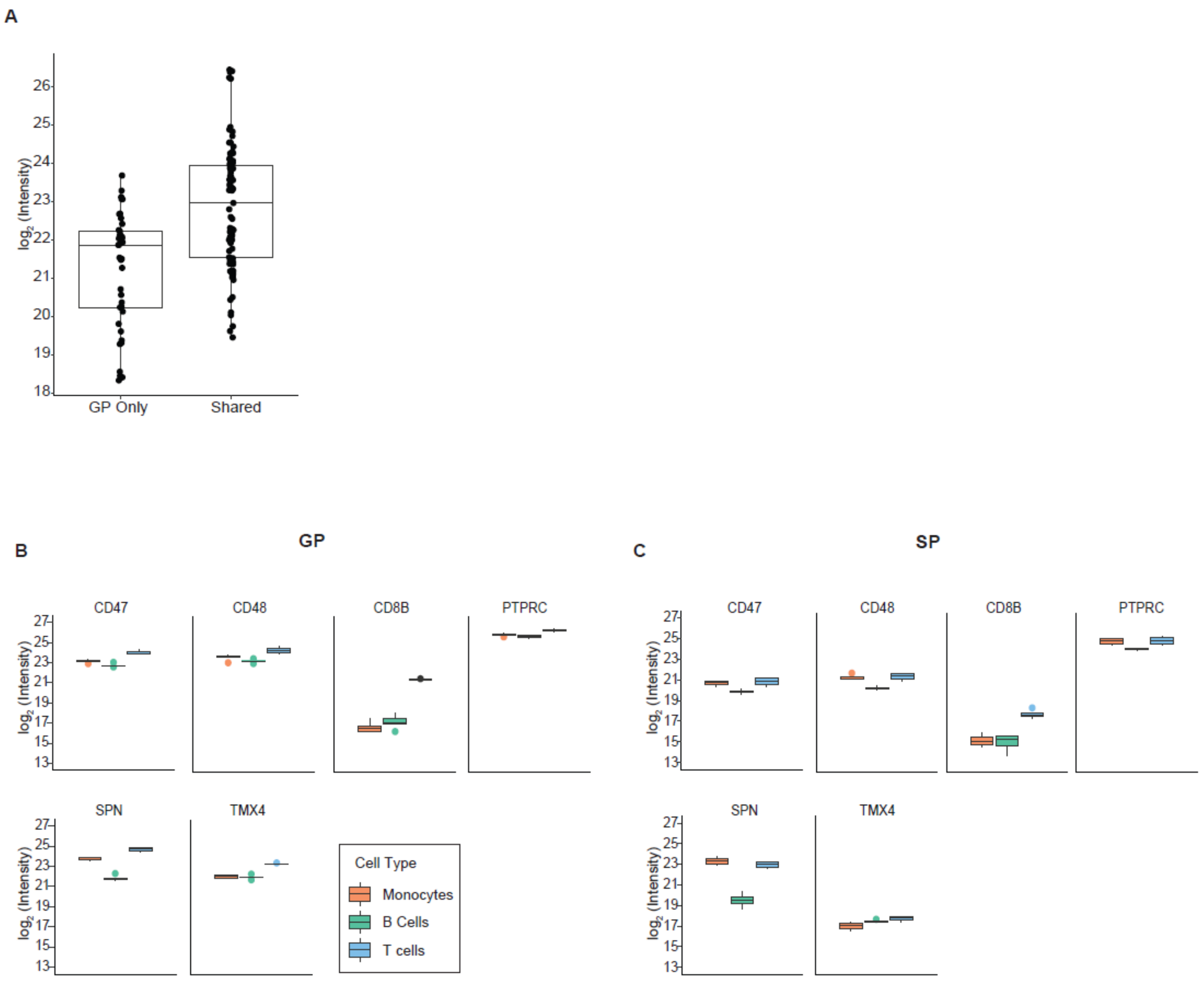

**Supplementary Figure S8**. Comparison of T-cell specific CSPs identified from SP and GP. (**A**) Boxplot for the log2 protein intensity of T-Cell enriched CSPs identified by either GP only or identified by both GP and SP (Shared). (**B**) Boxplot for the log2 GP protein intensity and (**C**) the log2 SP protein intensity of the 6 proteins that were T-Cell enriched by GP but not by SP.

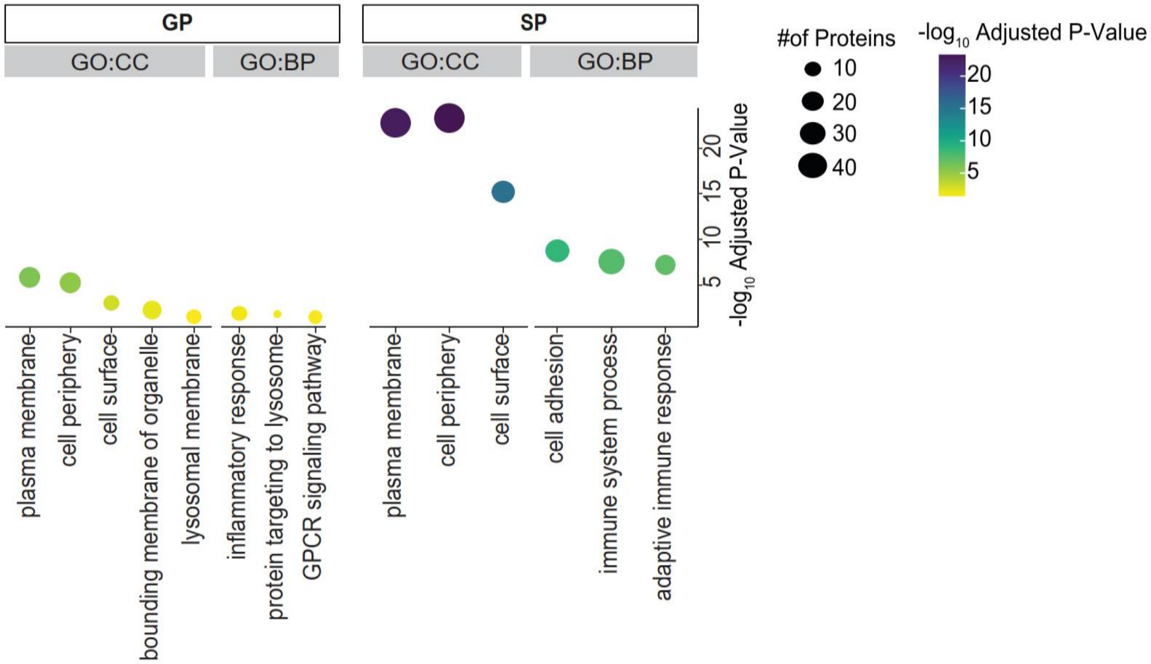

**Supplementary Figure S9**. Gene ontology of monocyte enriched CSPs from the SP workflow and the GP workflow.

All CSPs that were enriched in monocytes compared to B-Cells were selected for gene ontology analysis. Dotplot for the top enriched biological processes (GO:BP) and (GO:CC) for the SP and GP workflows are depicted. Size of the circles represent the number of proteins and the color of the circles represents the -log10 adjusted p-value of the enrichment.

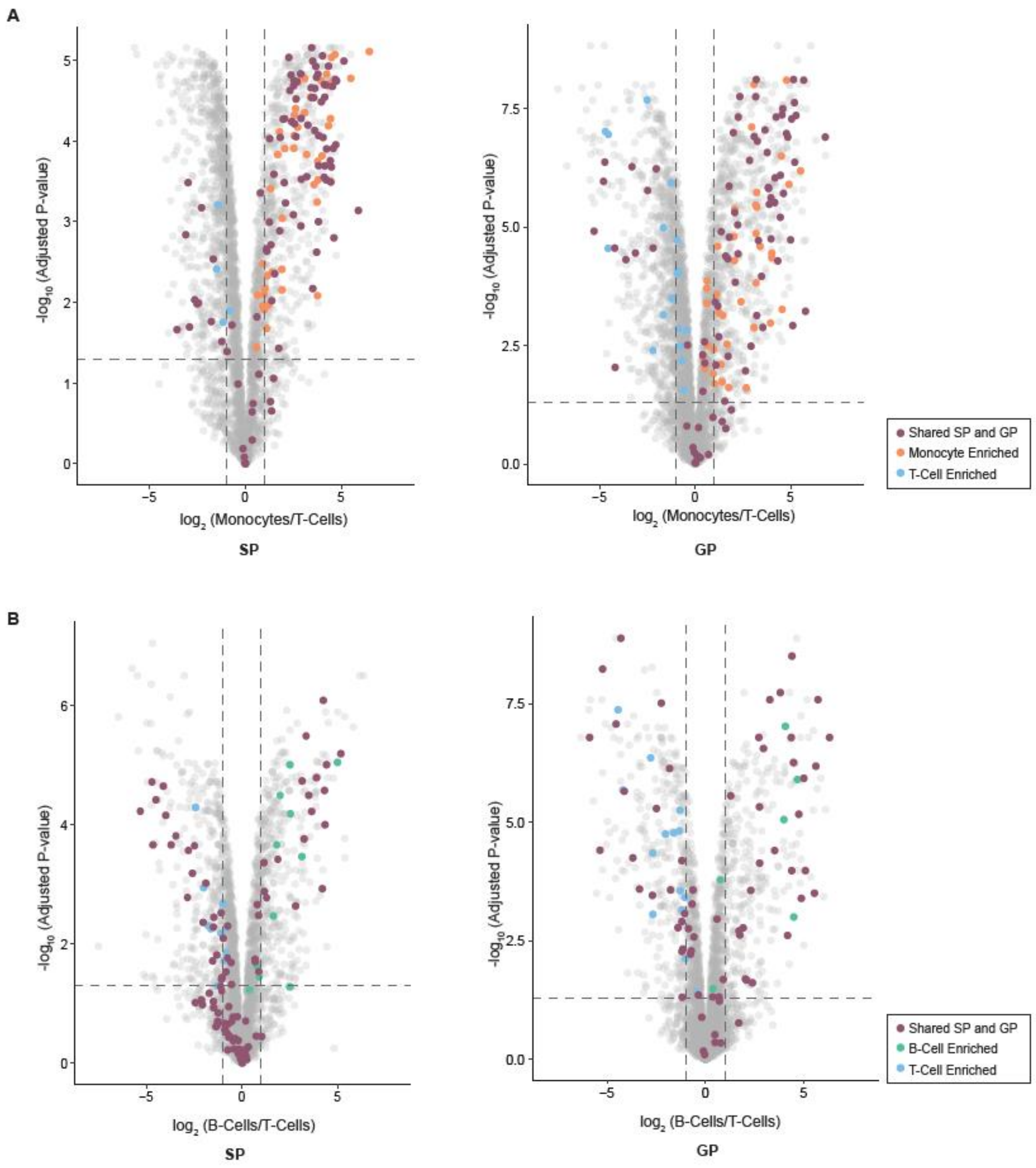

**Supplementary Figure S10**. Comparison of SP and GP for identifying cell-type-specific CSPs. (**A**) All proteins were tested for significance between monocytes and T-Cells from SP and GP workflows. Significant proteins in the volcano plot are defined by a P-value of <0.05 and fold change of >2. (**B**) All proteins were tested for significance between B-Cells and T-Cells from SP and GP workflows. Significant proteins in the volcano plot are defined by a P-value of <0.05 and fold change of >2.
